# Distinct Motor Cortex Somatotopy in Experimental Alzheimer’s Disease

**DOI:** 10.64898/2026.08.08.743539

**Authors:** SE Moss, CC Wolsh, RMII Brown, AR Brown, S Manchikalapudi, DQ Beversdorf, L Ma, JA Boychuk

## Abstract

Alzheimer’s Disease (AD) and related dementias (AD/RDs) impact cortical motor and sensory biology whereas the precise changes to these systems, and their clinical relevance, remain under debate. We hypothesized that cortical representations of complex and simple movements are differently altered during disease progression in 5XFAD mice, a well-established model of AD. Motor cortex somatotopy was determined in 5XFAD and Wild-Type Control (WT Control) mice at 6 and 12 months (mos.) of age using long-duration intracortical microstimulation (LD-ICMS) to systematically identify cortical sites evoking complex and simple forelimb movements. At 6 mos. of age, 5XFAD mice exhibited a significant expansion of motor cortical sites representing simple movements, specifically Elbow Flexion (p=0.0004) and Wrist Flexion (p=0.024). The over-sized territory for Elbow Flexion significantly distinguished 5XFAD from WT mice (Receiver Operating Characteristic [ROC] area under the curve [AUC]= 0.94, p= 0.0009) whereas discriminative performance of Wrist Flexion was a non-significant trend (AUC=0.75, p=0.059). By 12 mos. of age, motor cortex organization was markedly reorganized in 5XFAD mice, with significantly fewer cortical sites evoking complex Advance movement (p<0.0001) as well as simple Shoulder (p=0.0001), Elbow Extension (p=0.024), and Wrist Extension (p=0.003) movements. The number of sites for simple Wrist Flexion was significantly increased (p=0.011) in 12 mo. old 5XFAD mice. At 12 mos., territory size of several of these movement zones highly distinguished 5XFAD from WT mice, including Advance (AUC= 0.96, p= 0.0005), Shoulder (AUC= 0.97, p= 0.0004), Elbow Extension (AUC=0.78, p=0.034), Wrist Extension (AUC=0.85, p= 0.0082), and Wrist Flexion (AUC=0.80, p= 0.023). These findings demonstrate progressive, age-dependent remodeling of motor cortex somatotopy in 5XFAD mice, characterized by early expansion of specific simple movement cortical sites followed by deterioration of both complex and simple motor cortical maps as disease advances. Motor cortex somatotopic remodeling may provide a sensitive biomarker of AD/RDs progression.

## INTRODUCTION

Alzheimer’s Disease (AD) and related dementias (AD/RDs) are devastating disorders that create a massive economic burden in the United States and worldwide [1–4]. New research continues to improve interventions for AD/RDs whereas their implementation is hindered by the lack of sensitive biomarkers detecting the disease during its early stages. Approximately 41 million cases of AD/RDs are undiagnosed globally and world-wide under-diagnosed rates of these disorders may exceed 75% [5]. Earlier diagnosis is therefore essential to maximize therapeutic intervention benefit. Given that sensory and motor changes are implicated in these disorders [6–12], the present study sought to identify whether distinct motor cortex system-level changes occur in AD in a manner that can distinguish affected from unaffected. Accordingly, this study tested the hypothesis that 5XFAD mice [13], as a model of AD, present age-dependent remodeling in motor cortex somatotopy for complex and simple movements at 6 and 12 mos. of age relative to wild-type controls (WT Controls). These time-points evaluated comparatively early and late phases, of disease progression in 5XFAD mice, to serve as proof of concept for further optimizing early motor testing for the detection of these disease states. Translationally, these ages broadly correspond to early adult and middle-aged stages of human aging, respectively, although direct mouse-to-human age conversion remains approximate [14].

Sensory and motor cortices are often considered late-stage or less-affected structures in AD [15]. Disease progression is commonly defined pathologically by the accumulation of amyloid or tau depositions as a direct mechanism of dysfunction. Indeed, the 5XFAD model follows this progressive timeline with subcortical structures showing primary susceptibility, for accumulation of pathological proteins, prior to their manifestation in cortical structures [13,16–18]. Experimentally, cortical cells in AD exhibit many cell anatomical and physiological disruptions that precede overt accumulation of pathological proteins [13,16,19–21]. These findings suggest that cortical structures are impacted earlier in AD progression, than overt pathological proteins depositions may predict, and support the need to further understand how these systems are impacted in AD at cell and system levels. Sensory and motor systems are intriguing diagnostic targets due to their direct assessment of neural function, ease of activation, universality of testing equipment, and distinctiveness of responses.

Volitional motor control in mammals is a complex process requiring sensory and motor coupling that is sensitive to perturbation [22,23]. Neocortical layer 5 pyramidal cells (L5PCs) are a key component of motor control in mammals as their cell networks provide final brain motor outputs to spinal cord via the CST (i.e. corticospinal cells) and other sensorimotor pathways [24–26]. L5PCs and their signaling may be selectively vulnerable in AD/RDs. At the cellular level, neocortical L5PCs in 9 mo. old 5XFAD mice exhibit anatomical disruption including loss of this cell population and disruption of their dendritic arbors [13]. The timing of neocortical disease progression in 5XFAD mice has been further identified by several laboratories. For example, caspase signaling and cell loss of neocortical L5PCs has been detected at 4-6 mos. of age in 5XFAD mice [21]. Together, neocortical layer 5 is essential for descending somatic motor control and there is cellular evidence of its selective vulnerability to disease progression in 5XFAD mice.

Synaptic disruption of neocortical cells in 5XFAD mice precedes overt loss of cells and wide-scale morphological disruption. Ex vivo confocal microscopy of the bi-transgenic cross between 5XFAD and Thy1-YFP (H line) mice detects reduced basal dendrite spine density in 6 mo. old 5XFAD-Thy1-YFP mice that is not observed at 2 or 4 mos. of age mice [20]. In vivo 2-photon imaging of neocortex in 2.75-3.5 mo. old 5XFAD-Thy1-YFP mice fails to detect significant changes in overall L5PC structure and morphology[20]. Instead, L5PCs of 2-3 mo. old 5XFAD-Thy1-YFP mice exhibit reduced amplitude and frequency of miniature excitatory postsynaptic potentials (mEPSCs) indicative of synaptic dysfunction prior to anatomical injury [19]. Neocortical L5PCs of 5XFAD-Thy1-YFP mice also exhibit reduced intrinsic excitability including hyperpolarized membrane potentials, increased rheostat threshold to fire action potentials, as well as a loss of spike timing dependent plasticity during an electrical stimulation paradigm directed at neocortical layer 2/3 [19]. While these studies point to cellular disruption in AD/RDs, potential motor and sensory system-level changes in these disease states remain understudied.

At a system-level, motor cortex is somatotopically organized into territories generating distinct complex (i.e. multiple joints or axial muscles) and simple (i.e. single joint or muscle) outputs, that can be mapped by long-duration intracortical microstimulation (LD-ICMS) [27]. Motor cortex somatotopy has been determined with LD-ICMS in non-human primates [27–33], squirrels [29], tree shrews [28], and rodents [34–39]. LD-ICMS stimulation-evoked motor outputs display the time-scale and appearance of awake-behaving somatic motor control and remain discreet and stable in healthy conditions [27,31,34,38]. This functional organization of motor cortex is highly sensitive to changes in experience, excitability, neuronal signaling, and injury among other factors [36,39,40]. In mice, the motor cortex produces 4 types of “complex” (multi-joint) forelimb movement during LD-ICMS: Advance, Retract, Elevate and Dig and produces 4 types of “simple” (single-joint) forelimb movement: Digit, Wrist, Elbow and Shoulder [35].

Here, motor cortices of 5XFAD and WT Control mice were tested with LD-ICMS for complex and simple forelimb sub-types, as well as for hindlimb responses or co-responses of forelimb+hindlimb, to systematically determine this structure’s somatotopy. 5XFAD and WT Controls were compared by each movement sub-type’s number of responsive site(s) and motor cortical location(s) based on XY coordinates in two-dimensional space. Interestingly, 5XFAD and WT Controls differed in their proportions and locations of simple and complex movement sub-types at the two ages in separate ways. Receiver operating characteristic (ROC) curve analyses identified several individual somatotopic properties with robust performance for distinguishing between 5XFAD and WT Controls. These findings support motor cortex disruption in AD/RDs and suggest that changes to this system may be sensitive biomarkers in early identification and assessment of these disease states.

## MATERIALS AND METHODS

### SUBJECTS

Experimentation was performed on male Wild-Type Control (WT Control; C57BL/6J background) and [B6SJL-Tg (APPSwFlLon, PSEN1*M146L*L286V)6799Vas / Mmjax, RRID:MMRRC_034840-JAX] (i.e., 5XFAD) mice [13] at 6 and 12 mos. of age. 5XFAD mice over-express 3 mutations of human amyloid beta (A4) precursor protein 695 (APP) and 2 mutations of presenilin 1 (PSEN1) that are associated with Familial Alzheimer’s Disease (FAD); kindly donated by Robert Vassar, Ph.D., Northwestern University. Animals were obtained from the Mutant Mouse Resource and Research Center (MMRRC) and Jackson Laboratory (Bar Harbor, ME) at 4-6 weeks of age and raised in the local vivarium thereafter. Mice were housed with same-sex littermates (3-5/cage) in clear polycarbonate cages with sawdust bedding and enrichment toys on a 12h light/dark cycle at the University of Texas Health Science Center at San Antonio (UTHSCSA) and the University of Missouri-Columbia (MU). Standard mouse chow and water were provided ad libitum. All experimental procedures were approved by appropriate institutional animal care and use committees (UTHSCSA protocol 20180100AR; MU protocol 43673).

### Electrophysiological Mapping

Standard LD-ICMS techniques developed for mice were used to generate motor maps of motor cortical caudal (CFA) and rostral (RFA) forelimb areas [35,41]. Mice were anaesthetized with ketamine hydrochloride (150 mg/kg, i.p.) and xylazine (10 mg/kg, i.p.) at 6 mos. of age (WT Control, 30.19 ± 0.24 g; 5XFAD, 28.31 ± 0.57 g) or 12 mos. of age (WT Control, 32.01 ± 1.3 g; 5XFAD, 29.01 ± 0.65 g). Animals were secured in a stereotaxic frame on an aluminum block (7.5 L x 2.5 W x 2.0 H cm) to elevate the torso and allow free range of forelimb movement. A feedback-controlled heating pad-maintained core body temperature at ∼37.5°C. Supplemental injections of either ketamine (25 mg/kg) or a mixture of ketamine (10 mg/kg) and xylazine (1 mg/kg) were given as required throughout surgery to maintain a constant level of anesthesia as determined by monitoring vibrissae whisking, breathing rate, and cutaneous reflexes in response to a gentle foot/tail pinch (Table 1). A craniotomy was performed over the right motor cortex and physiological saline heated to body temperature was used to cover the cranial window to prevent cortical desiccation. An image of the exposed portion of the brain was captured using a digital camera coupled to a stereomicroscope (Olympus SZ61; Waltham, MA) and displayed on a personal computer. A grid of 500 µm squares was then overlaid on the digital image using imaging software (CorelDRAW; Ottawa, ON) and was calibrated to bregma using sagittal and frontal suture intersection coordinates obtained prior to the craniotomy. Electrode penetrations were performed at the intersections of the grid lines and in the center of each square to give an interpenetration distance of 354 µm, except when located over a blood vessel in which case a penetration was not performed or was performed at the minimal safe distance relative to the vessel.

**Table 1.**
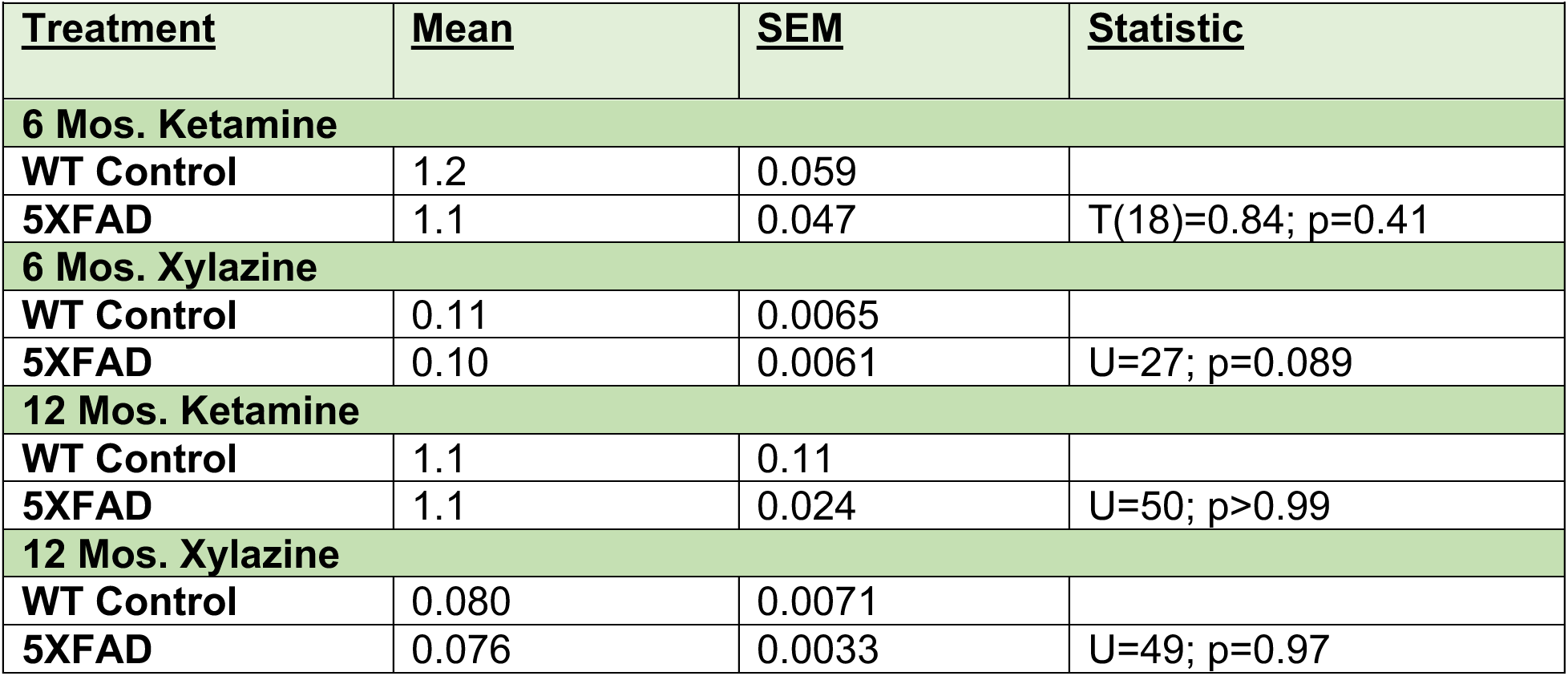
LD-ICMS Surgical Anesthetic (mg/hr).

| Table 1. LD-ICMS Surgical Anesthetic (mg/hr). |  |  |  |
| --- | --- | --- | --- |
| <u>Treatment</u> | <u>Mean</u> | <u>SEM</u> | <u>Statistic</u> |
| **6 Mos. Ketamine** |  |  |  |
| WT Control | 1.2 | 0.059 |  |
| 5XFAD | 1.1 | 0.047 | T(18)=0.84; p=0.41 |
| **6 Mos. Xylazine** |  |  |  |
| WT Control | 0.11 | 0.0065 |  |
| 5XFAD | 0.10 | 0.0061 | U=27; p=0.089 |
| **12 Mos. Ketamine** |  |  |  |
| WT Control | 1.1 | 0.11 |  |
| 5XFAD | 1.1 | 0.024 | U=50; p>0.99 |
| **12 Mos. Xylazine** |  |  |  |
| WT Control | 0.080 | 0.0071 |  |
| 5XFAD | 0.076 | 0.0033 | U=49; p=0.97 |

Platinum/iridium microelectrodes with an input impedance of 0.5 ± 0.1 MΩ (1000 Hz, 10 nA) were used (FHC Inc., Bowdoin, ME) were guided into the neocortex to a depth of 800 µm via microdrive (Narishige, Tokyo), corresponding to deep layer V, and adjusted to maximize the amplitude of evoked responses. Movements were readily elicited within a large depth range from surface (650-1000 µm) in mice with negligible effect on their nature or threshold [42]. An isolated pulse stimulator (Model 2100, A-M Systems; Carlsborg, WA) was used to deliver electrical current consisting of 500 ms trains of cathodal-leading 200 µs biphasic pulses delivered at a frequency of 333 Hz and an intensity of 100 µA consistent with previous studies [35]. Pulse trains were delivered in 0.2 Hz frequencies up to a maximum of 6 per site to ensure stability of evoked responses with stimulation sites evoking movement in ≥ 50% of trials considered responsive.

LD-ICMS testing and quantification was performed as described recently by the laboratory [35,41]. Mice were supported in a limb-free prone position with the wrist palm-down and digit, wrist, elbow, shoulder joints semi-flexed. Between stimulation trials, forelimb and hindlimb resting position was reset. Following the first stimulation site aimed at the center of the forelimb motor cortex, subsequent stimulations sampled in the parasagittal direction until either a non-limb or non-responsive point was observed. Mapping continued in a fashion to determine the border of the motor map first and then sampled toward the center on the overlaid grid until the border of the motor representation consisting of either a non-limb or non-responsive points was encountered. Motor map boundaries were defined by mapping all sites adjacent to a limb-responsive site as either non-responsive or non-limb. Throughout the surgery anesthetic levels were further monitored by verifying evoked movements in previously defined positive-response sites and surrounding non-responsive and non-forelimb points that defined the outside border of positive-response sites. Only complete LD-ICMS assessments wherein all forelimb-responsive sites and all hindlimb-responsive sites identified by these criteria were included in subsequent analyses.

Movements were monitored visually during electrophysiological mapping and video-recorded from a side view for offline assessment (100 frames/s, 2 ms shutter, Matrix Vision; Oppenwiler, Germany) with anatomical reference markers placed to identify the proximal termination of the humerus (shoulder), elbow, wrist, metacarpophalangeal joints, and the tip of the fourth digit with titanium dioxide paste to assist movement detection (Simi Motion; Unterschleißheim, Germany). A light*-*emitting diode synchronized with stimulator output was fixed to the stereotaxic frame in the camera field of view to determine stimulation onset. Evoked forelimb movements contralateral to the stimulated hemisphere were characterized as either complex when involving multiple joints during the stimulation or simple when involving a single joint (i.e., flexion/extension of the digit, wrist, elbow, or shoulder) according to previously defined criteria [35,41]. Forelimb movements that co-occurred with non-forelimb movements were classified as forelimb responses and the co-occurrence of hindlimb responses was recorded and quantified to determine all forelimb+hindlimb overlap. All hindlimb responses were tested and recorded according to these sampling parameters. The number and spatial location of responsive motor cortical sites for separate sub-types of movement were then calculated to perform group comparisons. For spatial properties, individual motor maps generated via LD-ICMS were spatially aligned, overlaid, and passed through a Gaussian smoothing filter to create group heatmaps in MATLAB R2024a (MathWorks, Natick, MA) based on adapted methodology [43].

### Statistical Analyses

Separate comparisons were made between WT Control and 5XFAD mice at 6 and 12 mos. of age. 1-Way Analysis of Variance (ANOVA) was used to compare amount administered anesthetic as well as number of somatotopic motor cortex sites and their within-group X or Y coordinates. 2-Way ANOVA was used to compare between-group X or Y coordinates for motor cortex sites using genotype and movement sub-type as factors. Post-hoc assessments were performed with unpaired t-tests, or Holm-Sidak procedure, as appropriate for number of multiple comparisons. Non-parametric tests (i.e., Mann-Whitney U) were performed for instances of unequal variance, based on Levene’s test and Bartlett’s test, or failure to meet assumptions of normality based on the D’Agostino-Pearson Omnibus K2 test. Receiver Operating Characteristic (ROC) area under the curve (AUC) values tested whether a randomly sampled 5XFAD mouse exhibited a more abnormal test score than a randomly chosen control; the corresponding p value tested the null hypothesis that AUC= 0.50 (no discrimination separate from chance). Statistical analyses were performed with GraphPad Prism 11 (GraphPad Software, La Jolla, CA). An experiment-wide α level of 0.05 was used and asterisks in figures indicate significance: \**p* ≤ 0.05, \*\**p* ≤ 0.01, \*\*\**p* ≤ 0.001. Reported *p-*values are adjusted for multiple comparisons. Data are reported as mean ± SEM with raw data points included in bar graphs.

## RESULTS

### Overall Motor Cortical Sites for *<u>Complex</u>* and *<u>Simple</u>* Movement Types

LD-ICMS was performed in WT Control and 5XFAD mice to define their motor cortex somatotopy of complex and simple movement sub-types. In all cases (1984 total test sites), the right hemisphere and left side of the body of each animal were analyzed. At 6 mos., a total of 891 electrode penetrations were performed in motor cortex of 20 mice (WT Control n=10; 5XFAD n=10). Responses comprised 249 forelimb, 158 non-forelimb, and 484 non-responsive stimulation sites. At 12 mos., a total of 1093 electrode penetrations were performed in motor cortex of 20 mice (WT Control n=10; 5XFAD n=10). Responses comprised 329 forelimb, 166 non-forelimb, and 598 non-responsive stimulation sites. The types of complex and simple forelimb movement were consistent with previous reports [35]. All forelimb movement sub-types were analyzed both for number of responsive sites as well as spatial topography based on XY cartesian coordinates. The number of responsive sites for hindlimb or the co-occurrence of hindlimb+forelimb were also determined. Comparisons of administered ketamine or xylazine failed to detect significant differences between 5XFAD and WT Control conditions at either age (Table 1).

The number of responsive sites in 6 mo. old WT Control and 5XFAD animals (Figure 1 A,B) were compared for all forelimb movements (Figure 1C), simple forelimb movements (Figure 1D), and complex forelimb movements (Figure 1E). In 6 mo. old 5XFAD animals, the number of responsive sites for all forelimb movements (15.29± 1.13) was not significantly different from WT Controls (15.71± 2.30), [U=24, p=0.15]. The number of responsive sites for complex forelimb movements did not significantly differ between 5XFAD and WT Control conditions at this age [T(18)=1.3; p=0.21]. In contrast, the number of responsive sites for simple forelimb movements was significantly increased in the 5XFAD group relative to WT Controls at 6 mos. of age [T(18)=2.7; p=0.02]. Mean values for number of responsive sites in WT Controls were as follows: complex forelimb (7.7± 0.45); simple forelimb (4.3± 0.58). Mean values for number of responsive sites in 5XFAD animals were as follows: complex forelimb (6.7± 0.63); simple forelimb (6.2± 0.42). Together, 5XFAD mice at 6 mos. fail to differ from WT Controls for number of all forelimb sites, or complex forelimb sites, whereas they exhibit an unexpected increase in the number of motor cortical sites for simple forelimb movements.

**Figure 1.**
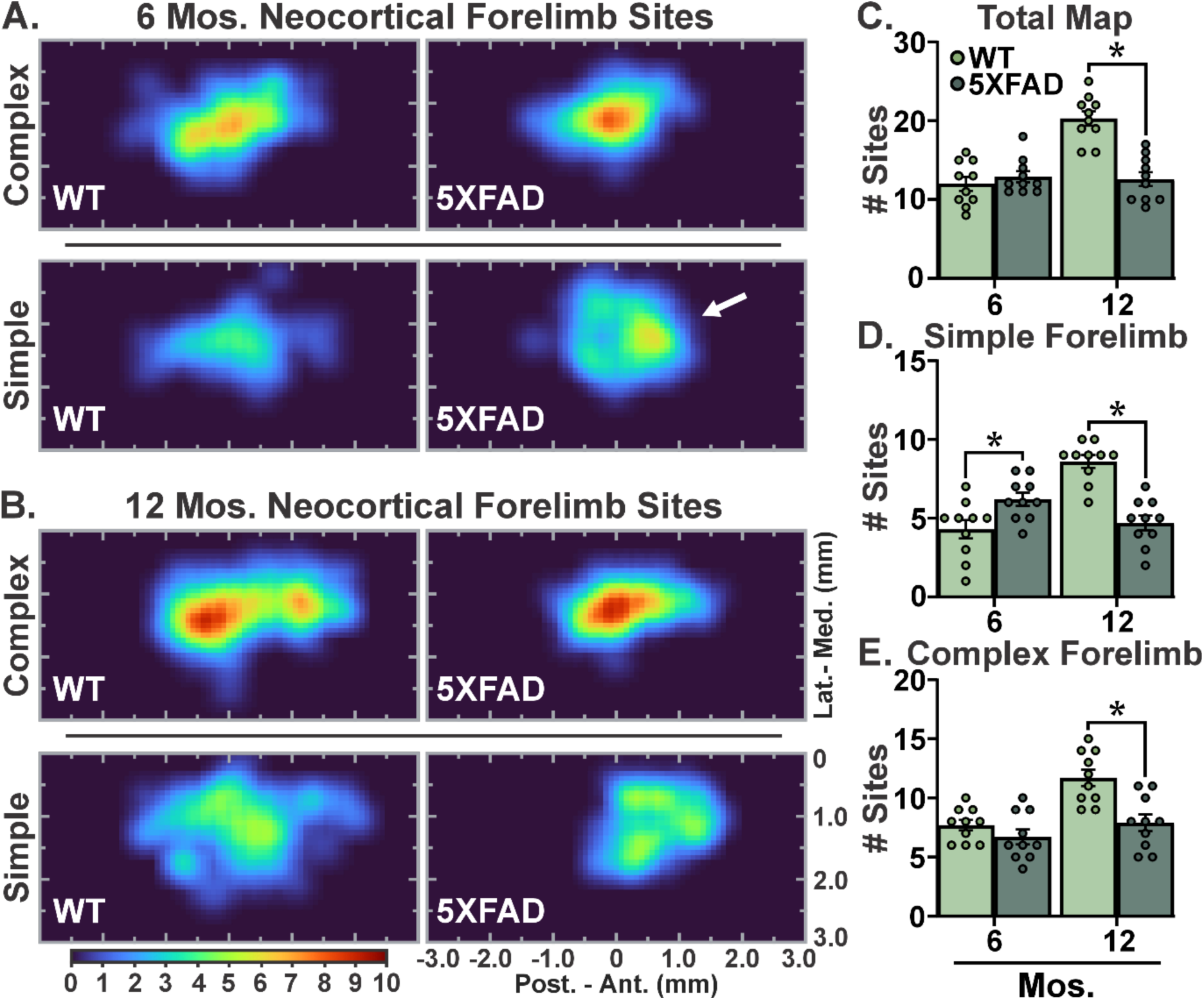
Separate Profiles of *<u>Overall</u>* Motor Cortex Somatotopy in 5XFAD and WT Control Mice. A-B. Representative heat map images of motor cortex somatotopy by group, age (6 and 12 mos.), and movement type (complex or simple). White arrow indicates distinct simple forelimb zone in 5XFAD mice. C. At 6 mos. of age there was no difference in the number of responsive sites for total forelimb movement between groups; in contrast, at 12 mos. of age the number of these sites was significantly reduced in the 5XFAD group relative to WT Controls. D. Unexpectedly, the number of responsive sites for simple forelimb movement was significantly greater in 5XFAD mice relative to WT Controls at 6 mos. of age whereas there were significantly fewer simple forelimb sites in 5XFAD mice at 12 mos. of age (p ≤ 0.05). E. The number of responsive sites for complex forelimb movement at 6 mos. of age was not affected in 5XFAD mice, relative to WT Controls, whereas there were significantly fewer complex forelimb sites in 5XFAD mice at 12 mos. of age (p ≤ 0.05). Bar graphs represent mean values± SEM with individual data points superimposed; All groups n=10; *= p ≤ 0.05.

The same somatotopic comparisons of movements (i.e., all forelimb, simple forelimb, and complex forelimb) were made between 5XFAD and WT Control groups at 12 mos. of age. In 12 mo. old animals, there were significantly fewer responsive sites for all forelimb movements in 5XFAD mice (14.00± 2.90) relative to WT Controls (24.71± 1.81), [U=6.5, p=0.0054]. The number of motor cortical sites for complex forelimb movements at 12 mos. of age was also significantly reduced in the 5XFAD group in comparison to WT Controls [T(18)=3.8; p=0.001]. Similarly, the number of motor cortical sites for simple forelimb movements at 12 mos. of age was significantly reduced in the 5XFAD group in comparison to WT Controls [T(18)=6.3; p<0.0001]. Mean values for number of responsive sites in WT Controls were as follows: complex forelimb (12± 0.70); simple forelimb (8.6± 0.40). Mean values for the number of responsive sites in 5XFAD animals were as follows: complex forelimb (7.9± 0.71); simple forelimb (4.7± 0.47). Thus, at 12 mos. of age, 5XFAD mice exhibited markedly fewer motor cortical sites for all forelimb movements, complex forelimb movements, and simple forelimb movements, suggesting broad progressive dysfunction in motor cortex’s capacity to produce somatic movement.

### <u>Six</u> Mos. of Age: <u>Number of Responsive Sites</u> for <u>Simple</u> Movement Sub-Types

Although a greater number of responsive sites for simple forelimb movement was detected in 5XFAD mice at 6 mos. (Figure 1D), the specific contribution of individual simple movement sub-types to this overall effect was strongly differential. The majority of simple movement sub-types did not significantly differ in number of responsive sites between groups at 6 mos. of age: Shoulder [U=46; p=0.77], Elbow extension [T(18)=0.97; p=0.34], Wrist Extension [T(18)=1.1; p=0.28], and Digit [T(18)=0.93; p=0.36] (Figure 2). In contrast, there was a significantly greater number of responsive sites for Elbow Flexion [U=6; p=0.0004] and Wrist Flexion [T(18)=2.5; p=0.024] in 5XFAD animals compared to WT Controls (Figure 2F). Mean values for number of responsive sites in WT Controls were as follows: Shoulder (1.2± 0.36); Elbow Flexion (0.3± 0.15); Elbow Extension (1.2± 0.33); Wrist Flexion (0.20± 0.13); Wrist Extension (0.50± 0.22); Digit (0.90± 0.23). Mean values for number of responsive sites in 5XFAD animals were as follows: Shoulder (1.3± 0.33); Elbow Flexion (1.9± 0.38); Elbow Extension (0.8± 0.25); Wrist Flexion (0.7± 0.15); Wrist Extension (0.9± 0.28); Digit (0.6± 0.22). Together, 6 mo. 5XFAD animals exhibited greater overall motor cortical territory for simple forelimb movement due in part to more responsive sites for simple elbow flexion and wrist flexion.

**Figure 2.**
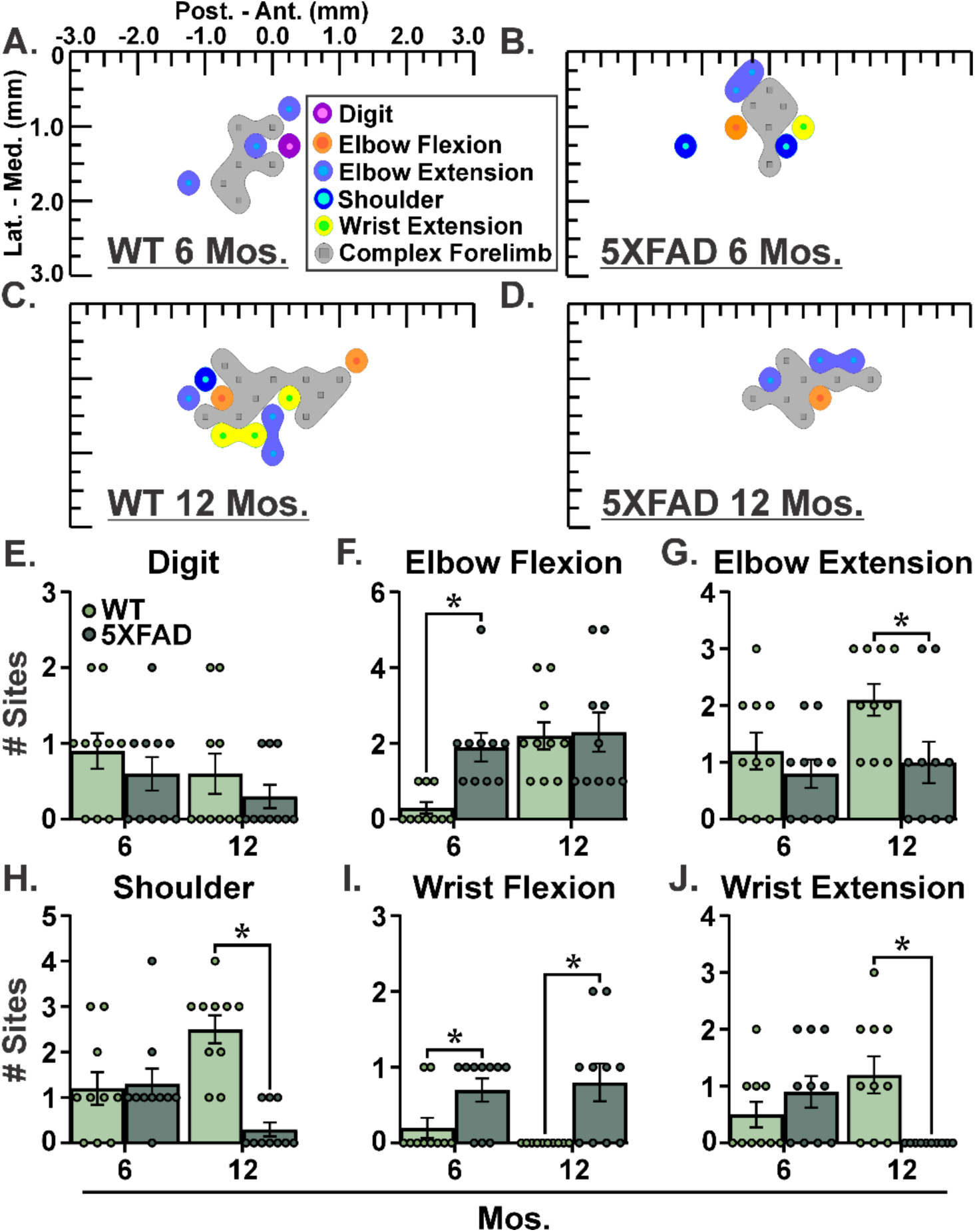
5XFAD Mice Exhibit Bi-directional Differences in Motor Cortex Somatotopy for *<u>Simple</u>* Movement Sub-types. A-D. Representative motor cortex somatotopy for simple movement sub-types in 5XFAD and WT Control groups at 6 and 12 mos. of age. E-J. At 6 mos. of age, 5XFAD and WT Control groups do not differ in number of responsive sites for the following simple movement sub-types: Digit, Wrist Extension, Elbow Extension, and Shoulder. Unexpectedly, 6 mo. old 5XFAD mice exhibit a significantly greater number of responsive sites for simple Elbow Flexion and Wrist Flexion (p ≤ 0.05). At 12 mos. of age, 5XFAD and WT Control groups do not differ in number of responsive sites for simple Digit and Elbow Flexion whereas they exhibit significantly fewer responsive sites for simple Shoulder, Elbow Extension, and Wrist Extension (p ≤ 0.05). Consistent with the earlier time-point, 12 mo. old 5XFAD mice exhibit a significantly greater number of responsive sites for simple Wrist Flexion (p ≤ 0.05). Bar graphs represent mean values± SEM with individual data points superimposed; All groups n=10; *= p ≤ 0.05.

### <u>Twelve</u> Mos. of Age: <u>Number of Responsive Sites</u> for <u>Simple</u> Movement Sub-Types

At 12 mos., total number of simple forelimb responsive sites in the 5XFAD group was significantly fewer relative to the WT Control group (Figure 1D) whereas specific contributions of individual simple movement sub-types to this overall effect were again strongly differential (Figure 2). When analyzed as sub-types for simple movement, Shoulder [U=3; p=0.0001], Elbow Extension [T(18)=2.4; p=0.024], and Wrist Extension [U=15; p=0.003] had significantly fewer responsive sites in the 5XFAD group compared to the WT Control group. In 5XFAD mice, there was a significant increase in responsive sites for Wrist Flexion compared to the WT Control group [U=20; p=0.011]. All other simple movement sub-types were not significantly different between groups at 12 mos. of age: Elbow Flexion [T(18)=0.16; p=0.88] and Digit [T(18)=0.98; p=0.34] (Figure 2). Mean values for number of responsive sites in WT controls were as follows: Shoulder (2.5± 0.31); Elbow Flexion (2.2± 0.36); Elbow Extension (2.1± 0.28); Wrist Flexion (0.0± 0); Wrist Extension (1.2± 0.33); Digit (0.6± 0.27). Mean values for number of responsive sites in 5XFAD animals were as follows: Shoulder (0.3± 0.15); Elbow Flexion (2.3± 0.52); Elbow Extension (1.0± 0.37); Wrist Flexion (0.80± 0.25); Wrist Extension (0.0± 0); Digit (0.30± 0.15). These results demonstrate an overall reduction in motor cortical territory for simple forelimb movements in 5XFAD animals at 12 mos. of age due to differential motor cortex somatotopy for simple movement sub-types.

### <u>Six</u> Mos. of Age: <u>Number of Responsive Sites</u> for <u>Complex</u> Movement Sub-Types

At 6 mos., the number of responsive sites of complex forelimb movements was not significantly affected in the 5XFAD group (Figure 1E). Analysis of the sub-types of complex motor cortical movement revealed no significant group changes at 6 mos. of age (Figure 3) for the number of responsive sites of Advance [T(18)=0.42; p=0.68], Elevate [T(18)=0.9, p=0.38], Retract [T(18)=0.63; p=0.53], and Dig [T(18)=1.7; p=0.10]. Mean values for number of responsive sites in WT controls were as follows: Advance (1.7± 0.37); Elevate (1.3± 0.21); Retract (2.7± 0.3); Dig (2.0± 0.37). Mean values for number of responsive sites in 5XFAD animals were as follows: Advance (1.5± 0.31); Elevate (1.0± 0.26); Retract (3.0± 0.37); Dig (1.2± 0.29). Thus, 5XFAD animals’ lack of overall change in complex motor cortical output at 6 mos. of age was not due to bidirectional changes across individual complex movement sub-types.

**Figure 3.**
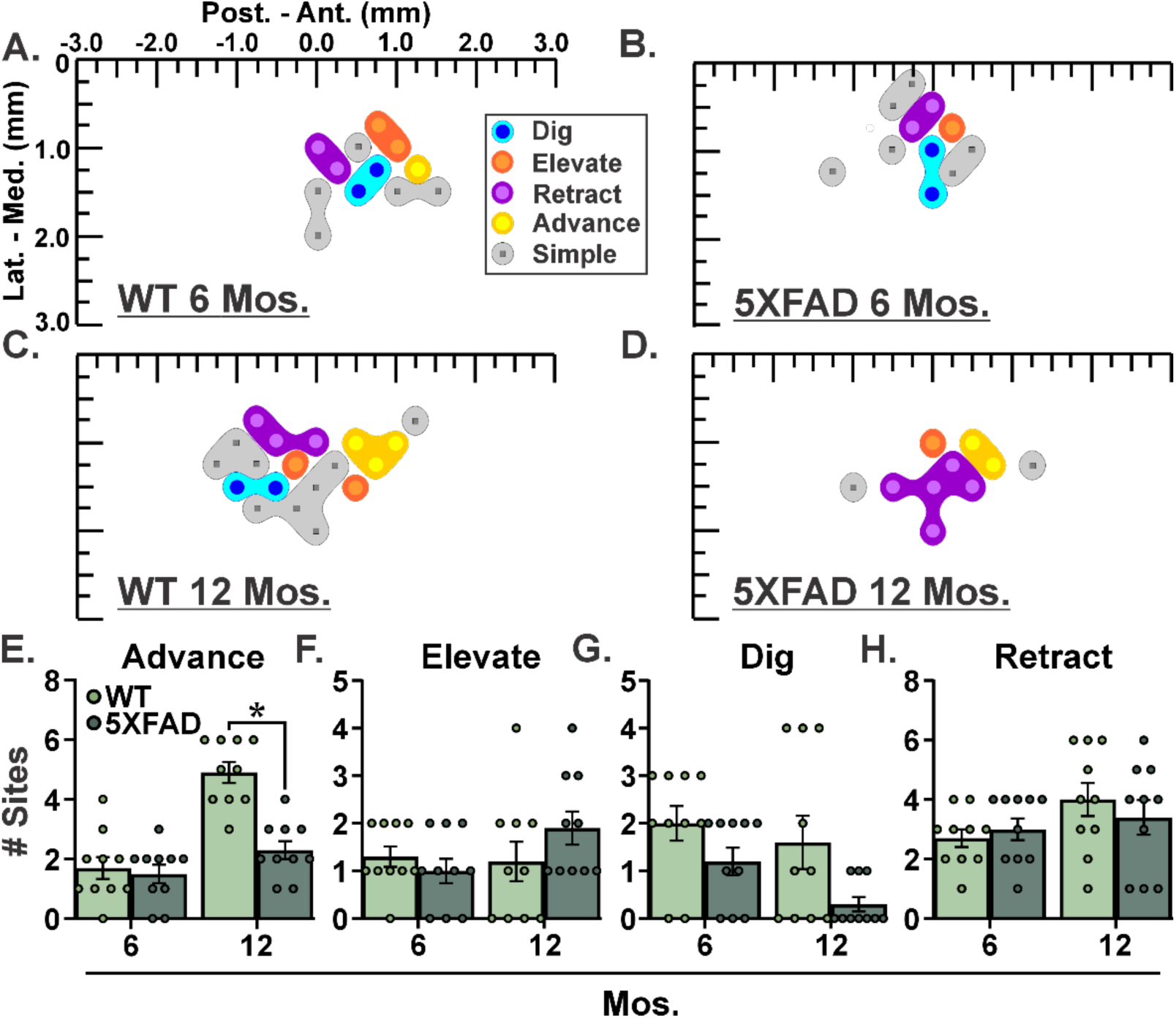
5XFAD Mice Exhibit Differential Motor Cortex Somatotopy for *<u>Complex</u>* Movement Sub-types. A-D. Representative motor cortex somatotopy for complex movement sub-types in 5XFAD and WT Control groups at 6 and 12 mos. of age. E-H. At 6 mos. of age, 5XFAD and WT Control groups do not significantly differ in number of responsive sites for any complex movement sub-types. At 12 mos. of age, 5XFAD and WT Control groups do not significantly differ in number of responsive sites for complex Elevate, Dig, and Retract whereas they exhibit significantly fewer responsive sites for complex Advance (p ≤ 0.05). Bar graphs represent mean values± SEM with individual data points superimposed; All groups n=10; *= p ≤ 0.05.

### Twelve Mos. of Age: Number of Responsive Sites for Complex Movement Sub-Types

Given significantly fewer responsive sites for total complex forelimb movements in the 5XFAD group at 12 mos. (Figure 1E), we then assessed the contribution of individual complex movement sub-types (Figure 3). Twelve mo. old 5XFAD mice exhibited significantly fewer sites of complex Advance [T(18)=5.7; p<0.0001] compared to the WT Control group. In contrast, the number of sites of Elevate [T(18)=1.3; p=0.21], Retract [T(18)=0.74; p=0.47] and Dig [U=29; p=0.10] were not significantly different between 5XFAD animals and WT Controls. Mean values for number of responsive sites in WT Controls were as follows: Advance (4.9± 0.35); Elevate (1.2± 0.42); Retract (4.0± 0.56); Dig (1.6±0.56). Mean values for number of responsive sites in 5XFAD animals were as follows: Advance (2.3± 0.30); Elevate (1.9± 0.35); Retract (3.4± 0.58); Dig (0.30± 0.15). Collectively, complex movement sub-types were differentially vulnerable in 5XFAD animals at 12 mos. of age despite an overall net reduction in the number of motor cortical sites for these movements.

#### Receiver Operating Characteristic (ROC) Curve Analyses of *<u>Number of Responsive Sites</u>* For Movement Sub-Types

The significant differences in number of responsive sites for several movement sub-types prompted us to test the discrimination performance of these properties for distinguishing WT Control and 5XFAD groups (Figure 4). All significant differences in number of responsive sites were subsequently scrutinized by ROC curve analysis. At 6 mos. of age, Elbow Flexion (AUC= 0.94, p= 0.0009, 95% confidence interval [CI]= 0.84 - 1.0) exhibited significant discrimination between WT Control and 5XFAD groups whereas Wrist Flexion exhibited a non-significant trend (AUC= 0.75, p= 0.059, 95% CI= 0.53 - 0.97). At 12 mos. of age the following simple movement sub-types exhibited significant discrimination between WT Control and 5XFAD groups: Shoulder (AUC= 0.97, p= 0.0004, 95% CI= 0.91 - 1.0), Elbow Extension (AUC= 0.78, p= 0.034, 95% CI= 0.56 - 1.0), Wrist Flexion (AUC= 0.80, p= 0.023, 95% CI= 0.59 - 1.0) and Wrist Extension (AUC= 0.85, p= 0.0082, 95% CI= 0.66 - 1.0). At 12 mos. of age, the complex sub-type Advance (AUC= 0.96, p= 0.0005, 95% CI= 0.88 - 1.0) exhibited significant discrimination between WT Control and 5XFAD groups. Collectively, the distinct number of responsive sites for several movement sub-types displayed robust significant discrimination performance between experimental AD/RDs and WT Controls.

**Figure 4.**
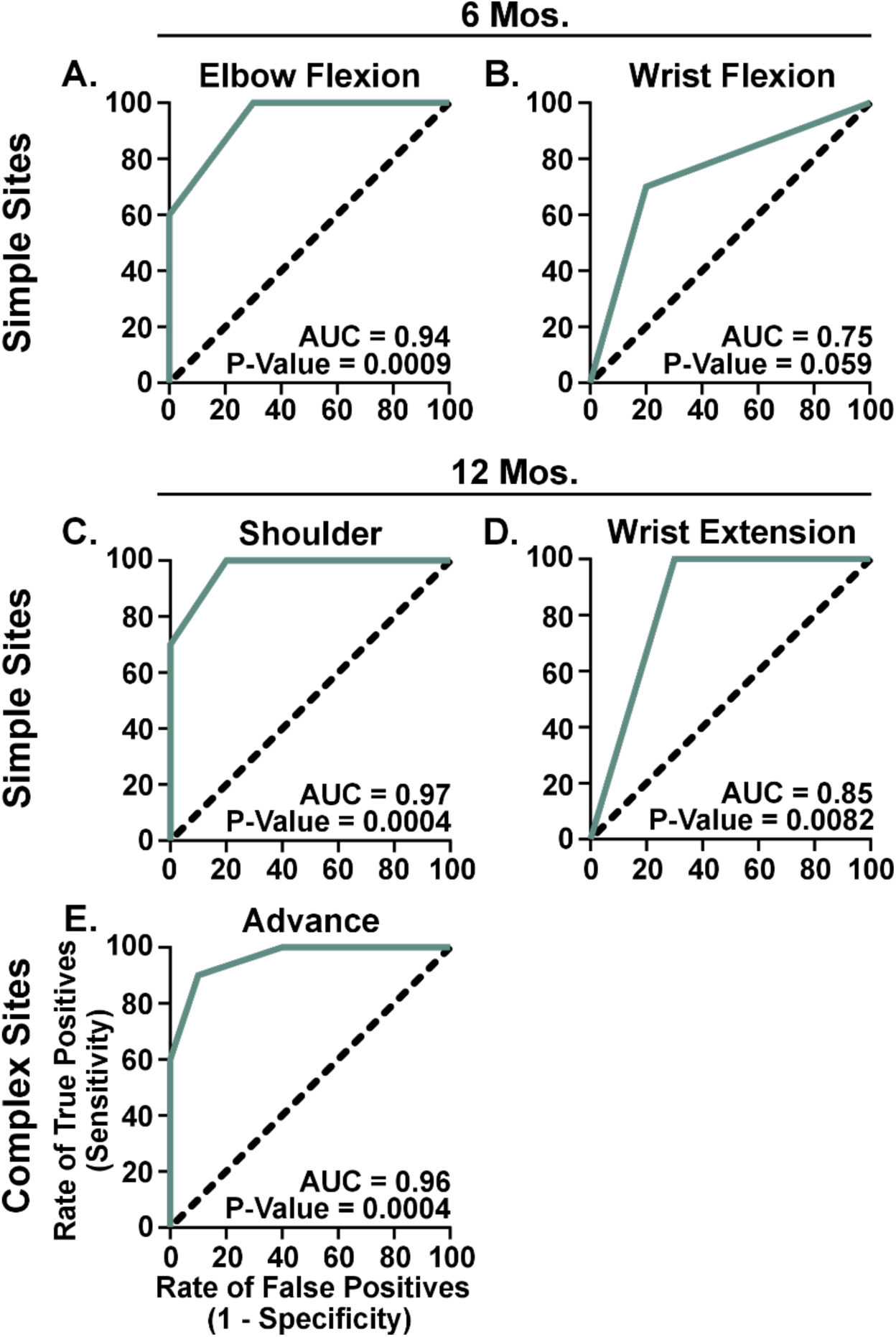
Motor Cortex Somatotopy of Movement Sub-types Distinguishes 5XFAD Mice From WT Controls. A-B. At 6 mos. of age, Receiver Operating Characteristic (ROC) curve analyses detect significant discrimination performance, between 5XFAD and WT Control groups, in the number of responsive sites for simple Elbow Flexion (p ≤ 0.05) and a statistical trend for simple Wrist Flexion (p= 0.059). C-E. At 12 mos. of age, ROC curve analyses detect significant discrimination performance, between 5XFAD and WT Control groups, in the number of responsive sites for simple Shoulder and Wrist Extension as well as in the number of responsive sites for complex Advance (p ≤ 0.05). AUC= Area Under the Curve.

### Overall *<u>Spatial Coordinates</u>* of *<u>Complex</u>* & *<u>Simple</u>* Motor Cortical Movements

Location of somatotopic zones is a key variable that can remodel in disease and injury [44–49]. To evaluate somatotopic location, forelimb and/or hindlimb responsive sites were assessed by their X and Y coordinates (mm) in relation to bregma (0,0) in 5XFAD and WT Control animals (Figures 5 & 6; Tables 2-6). Separate statistical analyses were performed for X and Y coordinates, 6- and 12-mo. ages, as well as simple and complex movement sub-types. At 6 mos. of age, there were no differences in location of the same movement sub-type in 5XFAD versus WT Control conditions (i.e., no main effects between groups) (Table 2). In contrast, 6 mo. old animals exhibited significant differences in location of separate movement sub-types, versus each other, within each group (i.e., significant main effects within-group; Table 2). At 12 mos. of age, a significant main effect between groups was detected for X coordinates of simple movements (p= 0.0042) whereas no other main effects of group were detected except for a statistical trend between groups for Y coordinates of complex movements (p= 0.053). As with 6 mos. of age, 12 mo. animals exhibited significant differences in location of separate movement sub-types, versus each other, within each group (Table 2). Together, there were few differences in location of the same somatotopic territories between 5XFAD versus WT Control conditions whereas each condition’s movement sub-types robustly differed from each other.

**Figure 5.**
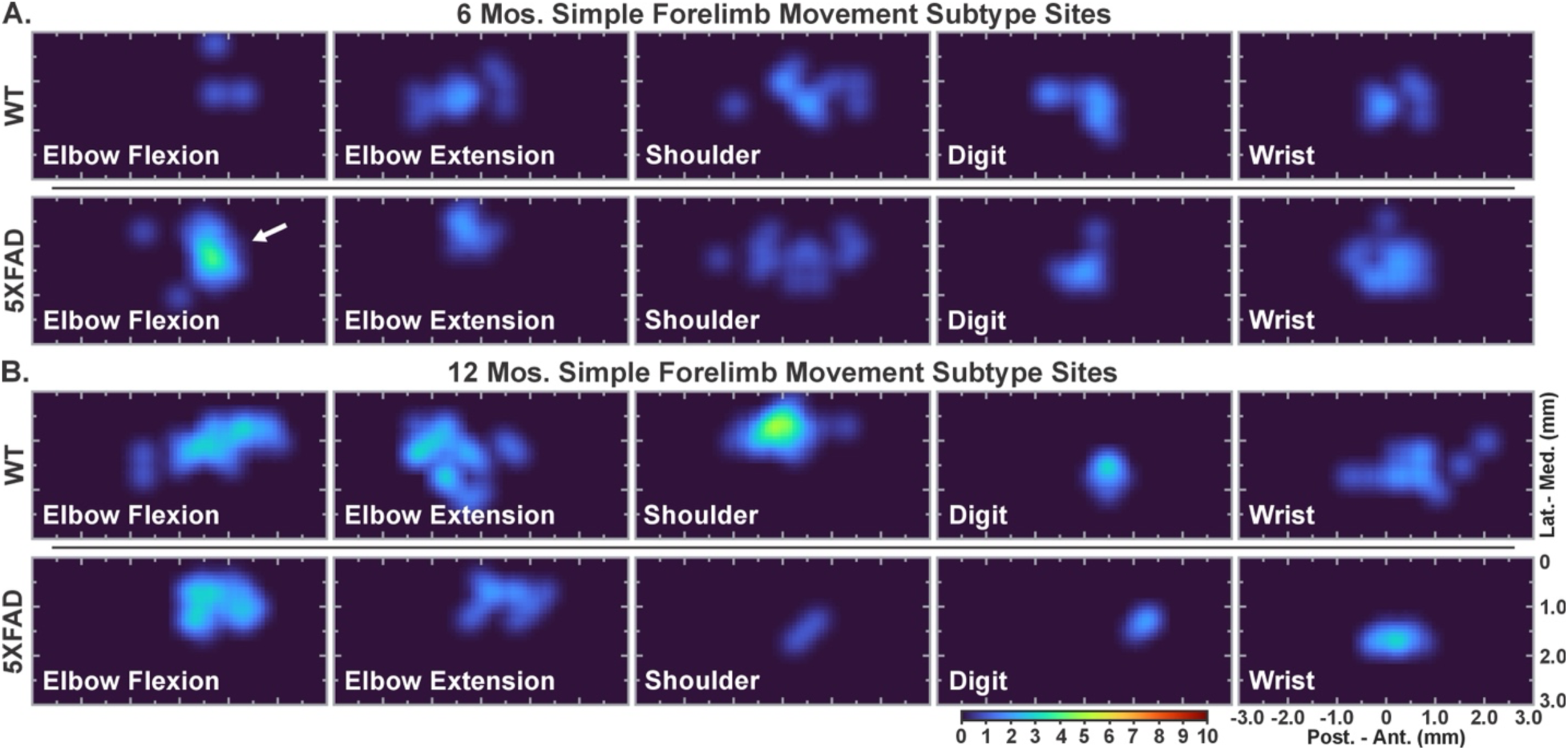
*<u>Simple</u>* Movement Sub-Type Differences in Spatial Distribution of Motor Cortex Somatotopy in 5XFAD and WT Control Mice. Representative heat map images of motor cortex somatotopy for simple movement sub-types for 5XFAD and WT Control conditions at 6 mos. (A) and 12 mos. of age (B). Few XY coordinates of individual simple movement sub-types differed between 5XFAD and WT Control conditions (Tables 2, 3 & 5). Instead, the majority of significant differences in XY coordinates were observed between separate simple movement sub-types within 5XFAD and WT Control conditions (Tables 2, 4 & 6). Collectively, somatotopic locations for simple movements are more likely to differ by sub-type than by experimental AD (i.e., 5XFAD). All groups n=10; *= p ≤ 0.05.

**Figure 6.**
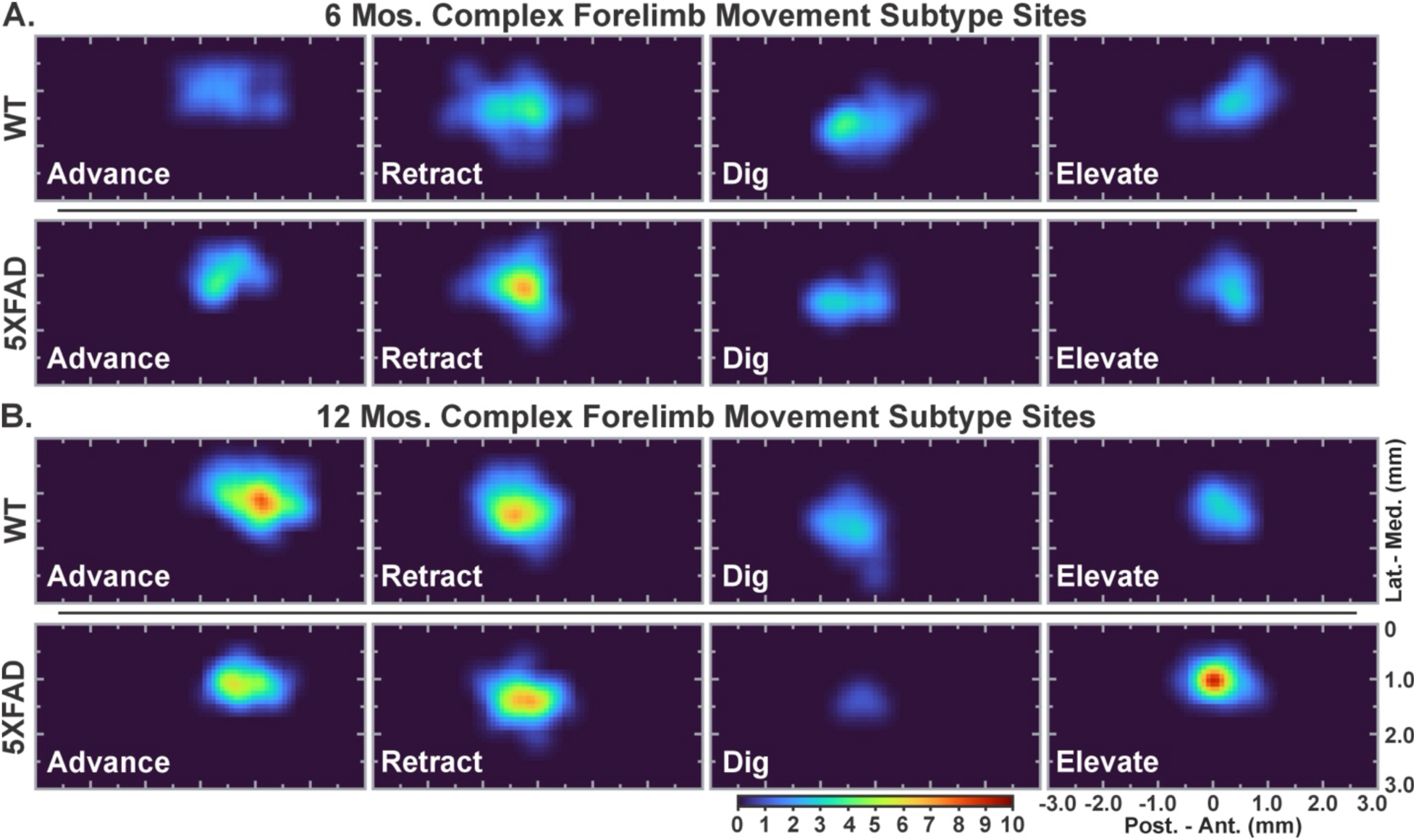
*<u>Complex</u>* Movement Sub-Type Differences in Spatial Distribution of Motor Cortex Somatotopy in 5XFAD and WT Control Mice. Representative heat map images of motor cortex somatotopy for complex movement sub-types for 5XFAD and WT Control conditions at 6 mos. (A) and 12 mos. of age (B). Few XY coordinates of individual complex movement sub-types differed between 5XFAD and WT Control conditions (Tables 2, 3 & 5). Instead, the majority of significant differences in XY coordinates were observed between separate complex movement sub-types within 5XFAD and WT Control conditions (Tables 2, 4 & 6). As with simple movements, somatotopic locations for complex movements are also more likely to differ by sub-type than by experimental AD (i.e., 5XFAD). All groups n=10; *= p ≤ 0.05.

**Table 2.** Main Effects of Overall Movement Type Spatial X & Y Coordinates (p values).

**6 Mos. Main Effects**
|  | **X-**<br>**Coordinates**<br>**(Simple)** | **Y-**<br>**Coordinates**<br>**(Simple)** | **X-**<br>**Coordinates**<br>**(Complex)** | **Y-**<br>**Coordinates**<br>**(Complex)** |
| --- | --- | --- | --- | --- |
| **Interaction** | 0.82 | *0.011 | 0.17 | 0.54 |
| **Genotype** | 0.28 | 0.18 | 0.26 | 0.18 |
| **Movement type** | ****<0.0001 | ***0.0002 | ****<0.0001 | ****0.0001 |

**12 Mos. Main Effects**
|  | **X-**<br>**Coordinates**<br>**(Simple)** | **Y-**<br>**Coordinates**<br>**(Simple)** | **X-**<br>**Coordinates**<br>**(Complex)** | **Y-**<br>**Coordinates**<br>**(Complex)** |
| --- | --- | --- | --- | --- |
| **Interaction** | ****<0.0001 | ***<0.0001 | *0.039 | 0.019 |
| **Genotype** | **0.0042 | 0.84 | >0.99 | 0.053 |
| **Movement type** | ****<0.0001 | ****<0.0001 | ****<0.0001 | ****0.0001 |

**Table 3.**
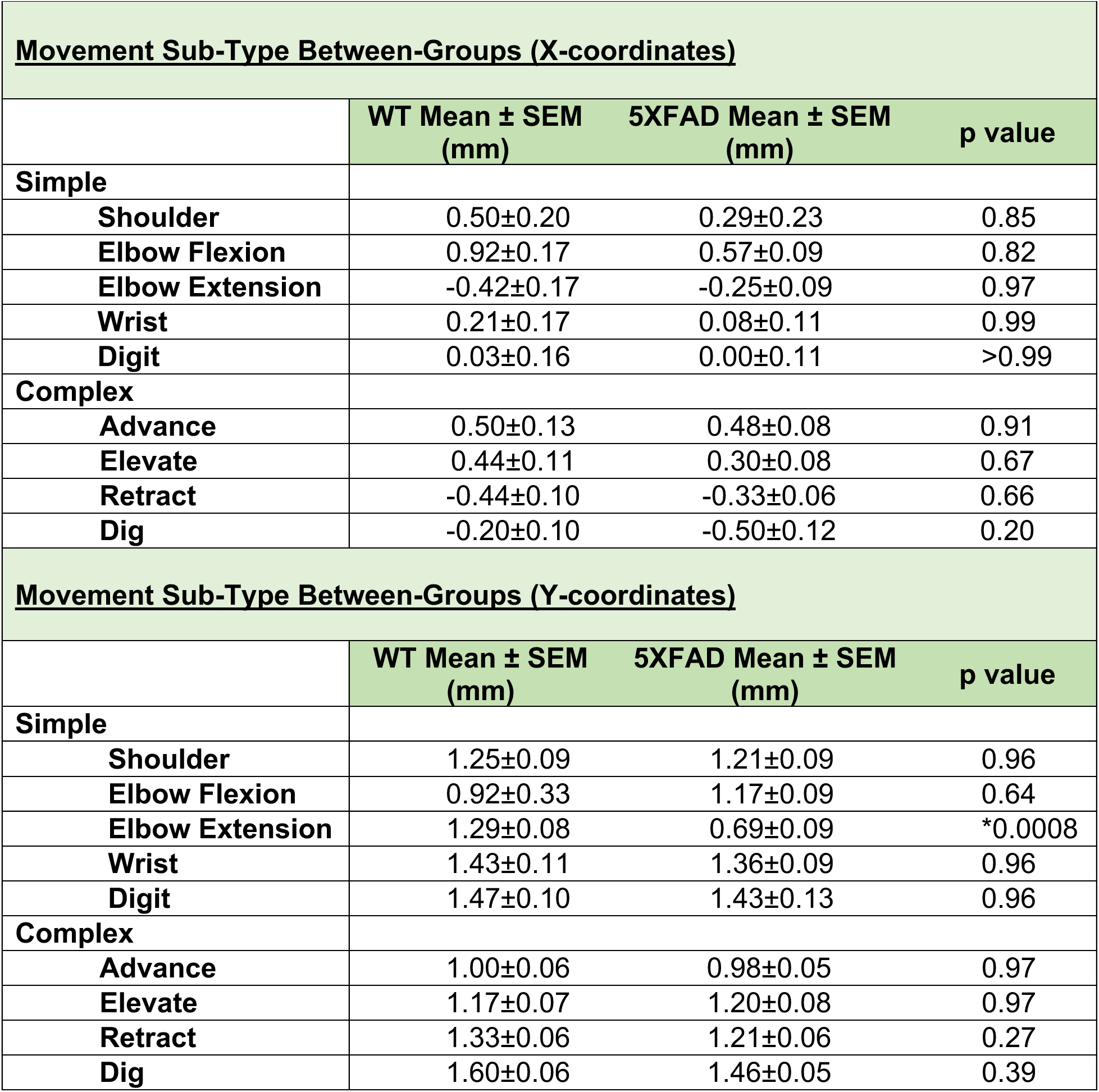
Between-Group Comparisons of <u>6 Mos.</u> X & Y Coordinates in Movement Sub-Types.

**Table 4.** Within-Group Comparisons of <u>6 Mos.</u> X & Y Coordinates in Movement Sub-Types (p values).

**6 Mos. Simple Movement Sub-Type Location Within Group (p values).**
| **WT** | **X-Coordinate Within Group** | **Y-Coordinate Within Group** |
| --- | --- | --- |
| Shoulder vs. Elbow Flexion | 0.61 | 0.54 |
| Shoulder vs. Elbow Extension | **0.004 | 0.93 |
| Shoulder vs. Wrist | 0.61 | 0.66 |
| Shoulder vs. Digit | 0.35 | 0.54 |
| Elbow Flexion vs. Elbow Extension | **0.008 | 0.44 |
| Elbow Flexion vs. Wrist | 0.36 | 0.18 |
| Elbow Flexion vs. Digit | 0.19 | 0.10 |
| Elbow Extension vs. Wrist | 0.19 | 0.74 |
| Elbow Extension vs. Digit | 0.36 | 0.66 |
| Wrist vs. Digit | 0.61 | 0.93 |
| **5XFAD** | **X-Coordinate Within Group** | **Y-Coordinate Within Group** |
| Shoulder vs. Elbow Flexion | 0.93 | 0.89 |
| Shoulder vs. Elbow Extension | 0.32 | *0.013 |
| Shoulder vs. Wrist | 0.99 | 0.60 |
| Shoulder vs. Digit | 0.93 | 0.58 |
| Elbow Flexion vs. Elbow Extension | ****<0.0001 | *0.013 |
| Elbow Flexion vs. Wrist | *0.02 | 0.48 |
| Elbow Flexion vs. Digit | *0.01 | 0.48 |
| Elbow Extension vs. Wrist | 0.28 | ***0.0004 |
| Elbow Extension vs. Digit | 0.60 | **0.0013 |
| Wrist vs. Digit | >0.99 | 0.89 |

**6 Mos. Complex Movement Sub-Type Location Within Group (p values).**
| **WT** | **X-Coordinate Within Group** | **Y-Coordinate Within Group** |
| --- | --- | --- |
| Advance vs. Elevate | 0.75 | 0.15 |
| Advance vs. Retract | ****<0.0001 | ***0.0008 |
| Advance vs. Dig | ***0.0002 | ****<0.0001 |
| Elevate vs. Retract | ****<0.0001 | 0.15 |
| Elevate vs. Dig | **0.0011 | ***0.0002 |
| Retract vs. Dig | 0.17 | **0.0050 |

| 5XFAD | X-Coordinate Within Group | Y-Coordinate Within Group |
| --- | --- | --- |
| Advance vs. Elevate | 0.27 | 0.1048 |
| Advance vs. Retract | ****<0.0001 | *0.045 |
| Advance vs. Dig | ****<0.0001 | ***0.0002 |
| Elevate vs. Retract | ****<0.0001 | 0.94 |
| Elevate vs. Dig | ****<0.0001 | 0.085 |
| Retract vs. Dig | 0.27 | *0.042 |

**Table 5.**
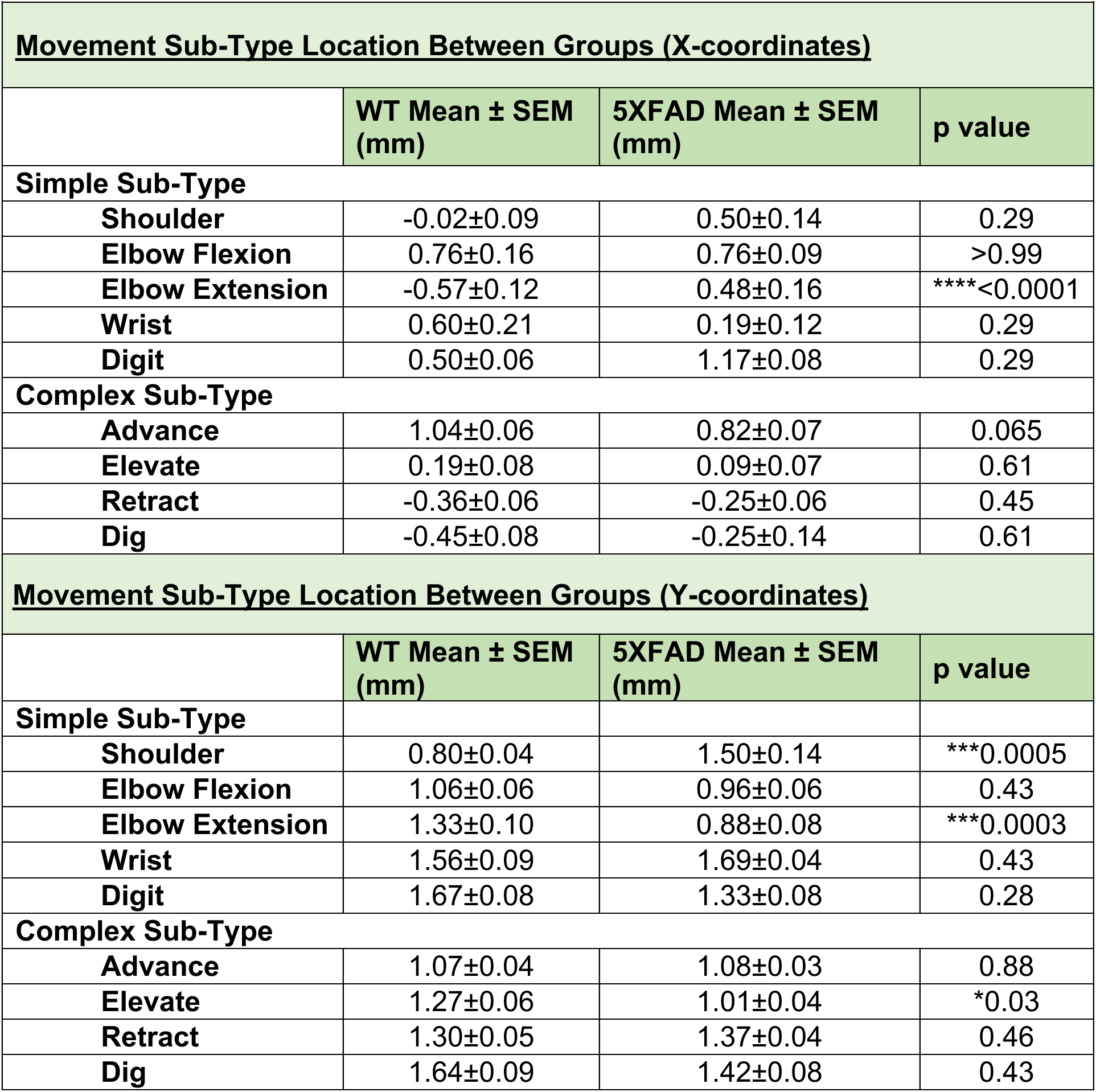
Between-Group Comparisons of <u>12 Mos.</u> X & Y Coordinates in Movement Sub-Types.

**Movement Sub-Type Location Between Groups (X-coordinates)**
|  | **WT Mean ± SEM**<br>**(mm)** | **5XFAD Mean ± SEM**<br>**(mm)** | **p value** |
| --- | --- | --- | --- |
| **Simple Sub-Type** |  |  |  |
| **Shoulder** | -0.02±0.09 | 0.50±0.14 | 0.29 |
| **Elbow Flexion** | 0.76±0.16 | 0.76±0.09 | >0.99 |
| **Elbow Extension** | -0.57±0.12 | 0.48±0.16 | ****<0.0001 |
| **Wrist** | 0.60±0.21 | 0.19±0.12 | 0.29 |
| **Digit** | 0.50±0.06 | 1.17±0.08 | 0.29 |
| **Complex Sub-Type** |  |  |  |
| **Advance** | 1.04±0.06 | 0.82±0.07 | 0.065 |
| **Elevate** | 0.19±0.08 | 0.09±0.07 | 0.61 |
| **Retract** | -0.36±0.06 | -0.25±0.06 | 0.45 |
| **Dig** | -0.45±0.08 | -0.25±0.14 | 0.61 |

**Movement Sub-Type Location Between Groups (Y-coordinates)**
|  | **WT Mean ± SEM**<br>**(mm)** | **5XFAD Mean ± SEM**<br>**(mm)** | **p value** |
| --- | --- | --- | --- |
| **Simple Sub-Type** |  |  |  |
| **Shoulder** | 0.80±0.04 | 1.50±0.14 | ***0.0005 |
| **Elbow Flexion** | 1.06±0.06 | 0.96±0.06 | 0.43 |
| **Elbow Extension** | 1.33±0.10 | 0.88±0.08 | ***0.0003 |
| **Wrist** | 1.56±0.09 | 1.69±0.04 | 0.43 |
| **Digit** | 1.67±0.08 | 1.33±0.08 | 0.28 |
| **Complex Sub-Type** |  |  |  |
| **Advance** | 1.07±0.04 | 1.08±0.03 | 0.88 |
| **Elevate** | 1.27±0.06 | 1.01±0.04 | *0.03 |
| **Retract** | 1.30±0.05 | 1.37±0.04 | 0.46 |
| **Dig** | 1.64±0.09 | 1.42±0.08 | 0.43 |

**Table 6.** Within-Group Comparisons of <u>12 Mos.</u> X & Y Coordinates in Movement Sub-Types (p values).

**12 Mos. Simple Movement Sub-Type Location Within Group (p values).**
| **WT** | **X-Coordinate Within Group** | **Y-Coordinate Within Group** |
| --- | --- | --- |
| Shoulder vs. Elbow Flexion | ****<0.0002 | *0.024 |
| Shoulder vs. Elbow Extension | *0.015 | ****<0.0001 |
| Shoulder vs. Wrist | *0.019 | ****<0.0001 |
| Shoulder vs. Digit | 0.22 | ****<0.0001 |
| Elbow Flexion vs. Elbow Extension | ****<0.0001 | 0.18 |
| Elbow Flexion vs. Wrist | 0.72 | ***0.0001 |
| Elbow Flexion vs. Digit | 0.72 | ***0.0003 |
| Elbow Extension vs. Wrist | ****<0.0001 | 0.089 |
| Elbow Extension vs. Digit | **0.0015 | 0.069 |
| Wrist vs. Digit | 0.73 | 0.51 |
| **5XFAD** | **X-Coordinate Within Group** | **Y-Coordinate Within Group** |
| Shoulder vs. Elbow Flexion | 0.62 | **0.004 |
| Shoulder vs. Elbow Extension | 0.93 | **0.002 |
| Shoulder vs. Wrist | 0.62 | 0.58 |
| Shoulder vs. Digit | 0.33 | 0.60 |
| Elbow Flexion vs. Elbow Extension | 0.38 | 0.60 |
| Elbow Flexion vs. Wrist | *0.015 | ****<0.0001 |
| Elbow Flexion vs. Digit | 0.47 | 0.065 |
| Elbow Extension vs. Wrist | 0.48 | ****<0.0001 |
| Elbow Extension vs. Digit | 0.12 | *0.032 |
| Wrist vs. Digit | *0.012 | 0.13 |

**12 Mos. Complex Movement Sub-Type Location Within Group (p values).**
| **WT** | **X-Coordinate Within Group** | **Y-Coordinate Within Group** |
| --- | --- | --- |
| Advance vs. Elevate | ****<0.0001 | 0.06 |
| Advance vs. Retract | ****<0.0001 | *0.0010 |
| Advance vs. Dig | ****<0.0001 | ****<0.0001 |
| Elevate vs. Retract | ****<0.0001 | 0.76 |
| Elevate vs. Dig | ***0.0001 | *0.0035 |
| Retract vs. Dig | 0.42 | *0.0007 |

| 5XFAD | X-Coordinate Within Group | Y-Coordinate Within Group |
| --- | --- | --- |
| Advance vs. Elevate | ****<0.0001 | 0.73 |
| Advance vs. Retract | ****<0.0001 | ****<0.0001 |
| Advance vs. Dig | ****<0.0001 | *0.11 |
| Elevate vs. Retract | **0.0013 | ****<0.0001 |
| Elevate vs. Dig | 0.18 | **0.0077 |
| Retract vs. Dig | >0.99 | 0.90 |

### *<u>Six</u>* Mos. of Age: *<u>Spatial Coordinates</u>* of *<u>Complex</u>* & *<u>Simple</u>* Motor Cortical Movement Sub-Types

To further examine group differences at 6 mos. of age, post hoc comparisons between 5XFAD versus WT Control groups detected a significant difference in Y coordinates of simple Elbow Extension (p=0.0008) whereas these groups failed to differ in all remaining comparisons regardless of group, movement type (simple or complex) and sub-type (Table 3). In contrast, many within-group comparisons were significant at 6 mos. in 5XFAD and WT Control groups; only a few of these effects occurred in simple movement sub-types whereas many differences were detected for complex movement sub-types (Table 4). At 6 mos., X coordinates for simple Elbow Flexion versus Elbow Extension were significantly different in both 5XFAD (p=0.008) and WT Controls (p<0.0001) groups while simple Shoulder versus Elbow Extension (p=0.004) was also significantly different in WT Controls (Table 4). Y coordinates for simple movement sub-type comparisons were only significantly different in the 5XFAD group at 6 mos. of age whereas the WT Control counterparts were non-significant (see Table 4 for complete list). Complex movement sub-types exhibited many differences from each other, for either X or Y coordinates, in both 5XFAD and WT Control groups (Table 4). Overall, 6 mos. somatotopic locations of movement sub-types did not substantially differ between experimental groups, whereas within each group the movement sub-types differed in location from each other, and these effects were most pronounced in complex movements rather than simple counterparts.

### *<u>Twelve</u>* Mos. of Age: *<u>Spatial Coordinates</u>* of *<u>Complex</u>* & *<u>Simple</u>* Motor Cortical Movement Sub-Types

At 12 mos. of age, a small number of significant differences were detected for the location of individual movement sub-types between 5XFAD and WT Control conditions (Table 5). 12 mo. old 5XFAD and WT Control groups significantly differed in simple Elbow Extension for X coordinates for (p<0.0001) and Y coordinates (p=0.0003; these groups also differed in Y coordinates for simple Shoulder (p=0.0005) and complex Elevate (p=0.03). As with 6 mos. ages, more significant differences were observed between movement sub-types within each experimental group at 12 mos. of age. In fact, WT Controls exhibited significant within-group differences in X coordinate comparisons of 6 simple and 5 complex movement sub-types (Table 6). This control condition also exhibited significant within-group differences in Y coordinate comparisons of 6 simple and 4 complex movement sub-types. 5XFAD animals exhibited significant within-group differences in X coordinate comparisons of 2 simple and 4 complex movement sub-types (Table 6). This experimental condition also exhibited significant within-group differences in Y coordinate comparisons of 5 simple and 4 complex movement sub-types. Collectively, there were few differences in the spatial location of simple and complex forelimb movements between 5XFAD and WT Control conditions, whereas in contrast, a larger number of significant effects were detected for how discrete motor outputs differ, in motor cortical location within group, and these effects were most prominent for complex rather than simple movement sub-types.

### *<u>Number of Responsive Sites</u>* for Hindlimb or Overlapping Forelimb+Hindlimb Movements

Our laboratory, and others, have previously observed positive correlations between behavioral recovery after brain insult and somatotopic zones that co-express forelimb and hindlimb movement or zones located at the motor cortical boundary between these movement types [39,41,50]. To address these possibilities in the context of AD and neurodegeneration, we quantified all hindlimb movement responses that shared a boundary with forelimb sites, or non-forelimb sites, as well as sites that co-expressed forelimb+hindlimb movement in 5XFAD mice and WT Controls either at 6 or 12 mos. of age (Figure 7). At 6 mos., the number of motor cortical hindlimb sites was not significantly different between groups [U=36, p=0.29] when comparing 5XFAD (mean: 6.40± 0.37) to WT Control (mean: 7.80± 0.83). To further test for a relationship between hindlimb and forelimb at 6 mos. of age, the numbers of motor cortical forelimb+hindlimb co-expressing sites were separately compared between groups where no significant difference was detected [U=50, p>0.99] between 5XFAD animals (mean: 3.70± 0.26) and WT Controls (mean: 4.10± 0.55). At 12 mos. of age, the number of motor cortical hindlimb sites was not significantly different between groups [U=44.5, p=0.69] when comparing 5XFAD (mean: 12.40± 0.87) to WT Controls (mean: 13.00± 0.71). The number of motor cortical forelimb+hindlimb co-expressing sites also exhibited no significance difference [T(18)=0.92; p=0.37] between 5XFAD animals (mean: 7.60± 0.58) and WT Controls (mean: 8.40± 0.65) at 12 mos. of age. Thus, 5XFAD conditions tended to exhibit slightly fewer hindlimb-related somatotopic sites numerically whereas these differences were not statistically significant.

**Figure 7.**
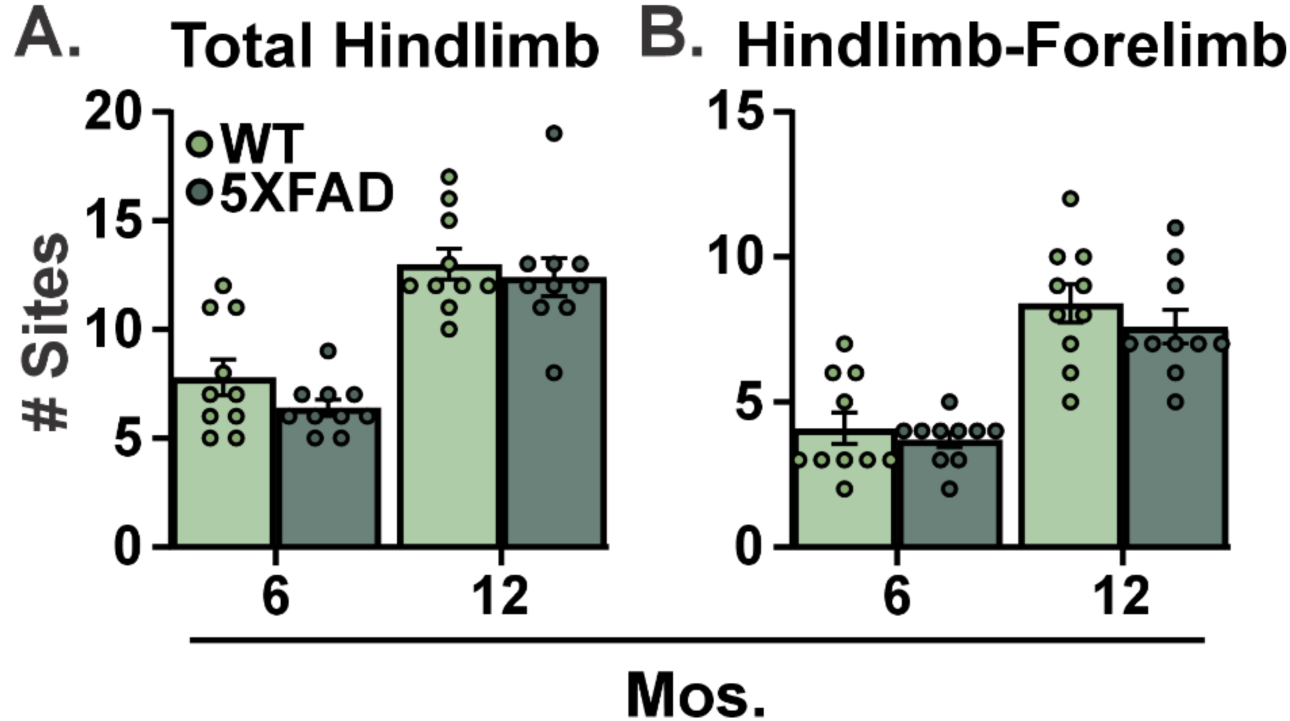
Unaffected Motor Cortex Somatotopy for Hindlimb and Dual Hindlimb-Forelimb Movements In 5XFAD Mice. A. At 6 and 12 mos. of age, 5XFAD and WT Control groups did not differ in the number of responsive sites for Hindlimb. B. At 6 and 12 mos. of age, 5XFAD and WT Control groups did not differ in the number of responsive sites co-expressing Hindlimb-Forelimb. Bar graphs represent mean values± SEM with individual data points superimposed; All groups n=10; *= p ≤ 0.05.

## DISCUSSION

Experiments here determined motor cortical somatotopy in 5XFAD and WT Control mice to understand whether this biology is uniquely impacted by Alzheimer’s Disease across separate ages. At 6 mos. of age, motor cortical somatotopy of these two groups was highly similar except for a notable increase in the motor cortex territory for producing simple movements in 5XFAD mice that was driven by select increase in specific simple movement sub-types. At 12 mos. of age 5XFAD mice exhibit fewer motor cortical sites capable of producing complex and simple movements without substantial changes in the spatial localization of discreet sub-types of movement (relative to WT Controls). Within each group, separate movement sub-types differed in spatial location from each other at 6 and 12 mos. of age. Importantly, motor cortex somatotopy in 5XFAD mice at 6 mos. was unique, compared to their 12 mo. counterparts, rather than merely a subtle early version of the later stage. Complex motor cortex outputs may rely more on synaptic integration of intracortical networks suggesting a breakdown of this biology in 12 mo. 5XFAD mice [38]. The present findings suggest that motor cortical somatotopy is distinct in experimental AD; a phenomenon that is dependent on both age and movement sub-type. These results support the possibility that functional organization of cortical systems may serve as a diagnostic feature in the early stages of AD.

At 6 mos. of age, the observed increase in motor cortical simple movements in 5XFAD mice may indicate biological dysfunction, compensation or be epiphenomenal. It is unlikely for shifts of motor cortical somatotopy to be epiphenomena given the large number of studies using ICMS in rodents wherein changes in the territories of complex and simple movement correlate with behavioral performance under healthy and pathological conditions [36,39,41,51–54]. For example, development of motor cortical somatotopy parallels motor performance [55,56]. Healthy motor forelimb training co-occurs with an increase in the number of motor cortical sites that co-express multiple and/or complex movement [36]. Limb immobility due to forelimb cast reduces the number of sites for motor cortical complex movement in a manner that is reversible [40]. Spared complex motor cortical output positively correlates with recovery from experimental brain insult [39,41,50]. Lesions or reversibly inactivation of identified regions of reorganized neocortex occludes rehabilitative-dependent gains in motor recovery while these administrations have limited effects in healthy control conditions [45–47,49].

At 12 mos., the dramatic loss of motor cortical sites for complex or simple movement in 5XFAD mice indicates a substantial diminishment of the somatic motor system’s capacity to produce volitional movement. Reduction of motor cortical sites to produce forelimb movement during LD-ICMS is associated with a loss of upper extremity function [36,39,41]. Interestingly however, both at 6 and 12 mos. of age the number of statistical differences for spatial coordinates of evoked motor responses between 5XFAD and WT Controls is sparse thereby suggesting a lack of change in central locations of these separate motor outputs across species, age and laboratory [34–36,39–41,43]. Instead, the 5XFAD condition exhibits a contraction of perimeter LD-ICMS forelimb responsive sites towards the center points of each movement type; this phenomenon is consistent with multiple manipulations tested by LD-ICMS including cortical cooling, experimental stroke injury, and peripheral limb immobility [36,40,41,57,58]. Together, the motor cortex somatotopy in 12 mos. 5XFAD mice suggests broad dysfunction rather than site-specific injury as is observed in acquired insults.

Motor cortical sites that co-express forelimb and hindlimb motor outputs, or exist at their somatotopic boundary, positively correlate with behavioral recovery after experimental stroke [41]; findings that have been previously reported in rats in separate acquired injury conditions [39,50]. Here, 5XFAD mice lacked overt change in motor cortical sites that express movement for hindlimb, or co-expression of forelimb+hindlimb, at both 6 and 12 mos. of age. This lack of change may represent an important feature of disease progression in AD. For example, there may be less possible remodeling of lower extremity somatotopic zones, to benefit upper extremity function, in AD compared to acquired injuries [39,41,50]. Additional lines of inquiry are needed to understand whether and how the somatotopic motor system remodels to sustain function in AD and neurodegenerative pathologies.

This study establishes several distinct features of motor cortex somatotopy in experimental AD disease progression. While motor cortex is generally considered a late structure to manifest gross anatomical disruption and accumulation of amyloid and tau in AD [15,59,60], other forms of early cellular disruption in these disease states remain possible. Accordingly, the 5XFAD model was selected to contextualize motor cortex somatotopy relative to known cellular changes in cortical and motor systems. At 4-6 mos., 5XFAD neocortex exhibits caspase signaling, anatomical disruption, and early cell loss of L5PCs [21]. Prior to 6 mos., synaptic dysfunction and intrinsic neurophysiological alterations occur in 5XFAD neocortex [19]. Thus, remodeling of motor cortex somatotopy in 6 mo. old 5XFAD mice may be driven by ongoing cell and anatomical damage as well as by the repercussions of early impairments with intrinsic and extrinsic cell physiology. The relationships between these biological events can be further resolved by testing motor cortex somatotopy at earlier ages of 5XFAD mice and by incorporating alterative models of AD. Substitute models of AD offer opportunities to focus on non-amyloid pathologies, non-APP and non-PSEN1 genes, alternate familial mutations, and knock-in versus over-expression systems, among other features.

Clinically, AD/RDs often present with cognitive impairment, memory loss, forms of aphasia, and confusion among other symptoms [61–63]. These behavioral impairments have guided intensive study of their corresponding neural systems including memory, speech and limbic structures [15,64]. Sensory and motor deficits are observed in AD/RDs [10] whereas debate remains for whether these behavioral changes are AD/RD-specific at early stages and whether comorbid neurological disorders play a role [6,15]. Mild cognitive impairment is associated with deficits in fine motor control while AD presents with deficits in fine and gross motor function [10]. Dementias are associated with reduced ability for grip strength and tactile sensation [11]. AD is linked to behavioral deficits in gait, based on one or several attributes, including slower walking speeds, reduced stride length, reduced axial balance, increased step variability and increased double support cycle time [7]. Disturbances in downstream neuromuscular junctions [65], or skeletal muscle [66,67], may contribute to motor deficits in AD/RDs including with gait.

For earlier detection, motor deficits in probable-mild AD are linked to impaired motor cortex, and descending motor control, as worsened pegboard dexterity and lower walking speed are associated with increased biomarker burden in motor cortex based on magnetic resonance imaging (MRI) and positron emission tomography (PET) targeting amyloid and tau [9]. Further, transcranial magnetic stimulation (TMS) of motor cortex using a rapid paired associative stimulation protocol fails to enhance motor evoked potentials in AD participants, or reduce their short afferent inhibition, despite successfully evoking these phenomena in controls [12]. TMS and theta burst stimulation of motor cortex have also identified moderate deficits in cortical plasticity, in mild cognitive impairment, that are more pronounced in prodromal AD and AD [8]. Consensus working groups highlight the ongoing need to identify optimal strategies for testing motor biology in AD/RDs using non-invasive and scalable technologies. Indeed, differences in the functional organization of motor cortex were recently found to distinguish obstructive sleep apnea patients, from health controls, based on focused strategies using MR neuroimaging [68].

In summary, 6 mo. old 5XFAD mice demonstrate noteworthy increases in the number of motor cortical sites to produce specific simple movement sub-types whereas somatotopic zones of other movement sub-types are unaffected or reduced. 5XFAD mice at 12 mos. demonstrate a more global reduction of motor cortical sites to produce both complex and simple movement that are combination of differential effects across movement sub-types. Changes in somatotopy of several specific movement sub-types robustly distinguished the 5XFAD condition from WT Controls. Spatially, there were few differences in the cartesian location of simple and complex forelimb movement sub-types between 5XFAD and WT Control conditions, whereas in contrast, a larger number of significant effects were detected for how discrete motor outputs differ in their motor cortical location within each experimental group; effects most prominent for complex rather than simple movement. These findings indicate that motor cortex organization and descending motor pathways are affected in experimental AD/RDs and support optimized testing of this biology for sensitive biomarkers of early detection and assessment of these disease states.

## Authors contributions

S.E.M, C.C.W., R.M.BII, A.R.B., and J.A.B conceived, designed and performed the experiments. S.E.M, C.C.W., R.M.BII, A.R.B., S.M and J.A.B analyzed data and prepared figures. All authors interpreted results of the experiments. All authors drafted the manuscript, revised it, and approved its final version.

## REFERENCES

[1] R. Brookmeyer, E. Johnson, K. Ziegler-Graham, H.M. Arrighi, Forecasting the global burden of Alzheimer’s disease, Alzheimer’s & dementia. 3 (2007) 186–191.

[2] A. Nandi, N. Counts, S. Chen, B. Seligman, D. Tortorice, D. Vigo, D.E. Bloom, Global and regional projections of the economic burden of Alzheimer’s disease and related dementias from 2019 to 2050: A value of statistical life approach, EClinicalMedicine. 51 (2022).

[3] D.P. Rice, P.J. Fox, W. Max, P.A. Webber, W.W. Hauck, D.A. Lindeman, E. Segura, The economic burden of Alzheimer’s disease care, Health affairs. 12 (1993) 164–176.

[4] C. Takizawa, P.L. Thompson, A. van Walsem, C. Faure, W.C. Maier, Epidemiological and economic burden of Alzheimer’s disease: a systematic literature review of data across Europe and the United States of America, Journal of Alzheimer’s disease. 43 (2014) 1271–1284.

[5] S. Gauthier, P. Rosa-Neto, J.A. Morais, C. Webster, World Alzheimer Report 2021: Journey through the diagnosis of dementia, Alzheimer’s Disease International. 2022 (2021) 30.

[6] M.W. Albers, G.C. Gilmore, J. Kaye, C. Murphy, A. Wingfield, D.A. Bennett, A.L. Boxer, A.S. Buchman, K.J. Cruickshanks, D.P. Devanand, At the interface of sensory and motor dysfunctions and Alzheimer’s disease, Alzheimer’s & Dementia. 11 (2015) 70–98.

[7] J. Andrade-Guerrero, H. Martínez-Orozco, M.M. Villegas-Rojas, A. Santiago-Balmaseda, K.M. Delgado-Minjares, I. Pérez-Segura, M.T. Baéz-Cortés, M.A. Del Toro-Colin, M. Guerra-Crespo, O. Arias-Carrión, Alzheimer’s disease: understanding motor impairments, Brain sciences. 14 (2024) 1054.

[8] F. Di Lorenzo, C. Motta, E.P. Casula, S. Bonnì, M. Assogna, C. Caltagirone, A. Martorana, G. Koch, LTP-like cortical plasticity predicts conversion to dementia in patients with memory impairment, Brain Stimulation. 13 (2020) 1175–1182.

[9] L. Gupta, Y. Ma, A. Kohli, K.L. Yang, J.M. Oh, T.J. Betthauser, N.A. Chin, O.C. Okonkwo, M.E. Pasquesi, V. Nair, Alzheimer’s disease biomarker burden in primary motor cortices is associated with poorer dexterity performance, Alzheimer’s & Dementia. 20 (2024) 5792–5799.

[10] A. Kluger, J.G. Gianutsos, J. Golomb, S.H. Ferris, A.E. George, E. Franssen, B. Reisberg, Patterns of motor impairment in normal aging, mild cognitive decline, and early Alzheimer’Disease, The Journals of Gerontology Series B: Psychological Sciences and Social Sciences. 52 (1997) P28–P39.

[11] E. Schonfeld, E. Schonfeld, C. Aman, N. Gill, D. Kim, S. Rabin, B. Shamshuddin, L. Sealey, R.G. Senno, Lateralized deficits in motor, sensory, and olfactory domains in dementia, Journal of Alzheimer’s Disease. 79 (2021) 1033–1040.

[12] C. Terranova, A. Sant’Angelo, F. Morgante, V. Rizzo, R. Allegra, M.G. Arena, L. Ricciardi, M.F. Ghilardi, P. Girlanda, A. Quartarone, Impairment of sensory-motor plasticity in mild Alzheimer’s disease, Brain stimulation. 6 (2013) 62–66.

[13] H. Oakley, S.L. Cole, S. Logan, E. Maus, P. Shao, J. Craft, A. Guillozet-Bongaarts, M. Ohno, J. Disterhoft, L. Van Eldik, R. Berry, R. Vassar, Intraneuronal β-amyloid aggregates, neurodegeneration, and neuron loss in transgenic mice with five familial Alzheimer’s disease mutations: potential factors in amyloid plaque formation, Journal of Neuroscience. 26 (2006) 10129–10140.

[14] S. Dutta, P. Sengupta, Men and mice: relating their ages, Life sciences. 152 (2016) 244–248.

[15] M.A. DeTure, D.W. Dickson, The neuropathological diagnosis of Alzheimer’s disease, Molecular neurodegeneration. 14 (2019) 32.

[16] S. Forner, S. Kawauchi, G. Balderrama-Gutierrez, E.A. Kramár, D.P. Matheos, J. Phan, D.I. Javonillo, K.M. Tran, E. Hingco, C. Da Cunha, Systematic phenotyping and characterization of the 5xFAD mouse model of Alzheimer’s disease, Scientific data. 8 (2021) 270.

[17] R. Gail Canter, W.-C. Huang, H. Choi, J. Wang, L. Ashley Watson, C.G. Yao, F. Abdurrob, S.M. Bousleiman, J.Z. Young, D.A. Bennett, 3D mapping reveals network-specific amyloid progression and subcortical susceptibility in mice, Communications biology. 2 (2019) 360.

[18] A.L. Oblak, P.B. Lin, K.P. Kotredes, R.S. Pandey, D. Garceau, H.M. Williams, A. Uyar, R. O’Rourke, S. O’Rourke, C. Ingraham, Comprehensive evaluation of the 5XFAD mouse model for preclinical testing applications: a MODEL-AD study, Frontiers in aging neuroscience. 13 (2021) 713726.

[19] Y. Buskila, S.E. Crowe, G.C. Ellis-Davies, Synaptic deficits in layer 5 neurons precede overt structural decay in 5xFAD mice, Neuroscience. 254 (2013) 152–159.

[20] S.E. Crowe, G.C. Ellis-Davies, Spine pruning in 5xFAD mice starts on basal dendrites of layer 5 pyramidal neurons, Brain Structure and Function. 219 (2014) 571–580.

[21] W.A. Eimer, R. Vassar, Neuron loss in the 5XFAD mouse model of Alzheimer’s disease correlates with intraneuronal Aβ 42 accumulation and caspase-3 activation, Molecular neurodegeneration. 8 (2013) 1–12.

[22] J. Merel, M. Botvinick, G. Wayne, Hierarchical motor control in mammals and machines, Nature communications. 10 (2019) 5489.

[23] S.H. Scott, Optimal feedback control and the neural basis of volitional motor control, Nature Reviews Neuroscience. 5 (2004) 532–545.

[24] H. Asanuma, H. Sakata, Functional organization of a cortical efferent system examined with focal depth stimulation in cats, J. Neurophysiol. 30 (1967) 35–54.

[25] H. Asanuma, J. Ward, Patterns of contraction of distal forelimb muscles produced by intracortical stimulation in cats, Brain research. 27 (1971) 97–109.

[26] S. Ghosh, R. Porter, Morphology of pyramidal neurones in monkey motor cortex and the synaptic actions of their intracortical axon collaterals, The Journal of physiology. 400 (1988) 593–615.

[27] M.S. Graziano, C.S. Taylor, T. Moore, Complex movements evoked by microstimulation of precentral cortex, Neuron. 34 (2002) 841–851.

[28] M.K. Baldwin, D.F. Cooke, L. Krubitzer, Intracortical microstimulation maps of motor, somatosensory, and posterior parietal cortex in tree shrews (Tupaia belangeri) reveal complex movement representations, Cerebral cortex. 27 (2016) bhv329.

[29] D.F. Cooke, J. Padberg, T. Zahner, L. Krubitzer, The functional organization and cortical connections of motor cortex in squirrels, Cerebral cortex. 22 (2012) 1959–1978.

[30] O.A. Gharbawie, I. Stepniewska, J.H. Kaas, Cortical connections of functional zones in posterior parietal cortex and frontal cortex motor regions in new world monkeys, Cerebral cortex. 21 (2011) 1981–2002.

[31] D.M. Griffin, H.M. Hudson, A. Belhaj-Saïf, P.D. Cheney, EMG Activation Patterns Associated with High Frequency, Long-Duration Intracortical Microstimulation of Primary Motor Cortex, The Journal of Neuroscience. 34 (2014) 1647–1656.

[32] A. Mayer, M.K. Baldwin, D.F. Cooke, B.R. Lima, J. Padberg, G. Lewenfus, J.G. Franca, L. Krubitzer, The multiple representations of complex digit movements in primary motor cortex form the building blocks for complex grip types in capuchin monkeys, Journal of Neuroscience. 39 (2019) 6684–6695.

[33] I. Stepniewska, O.A. Gharbawie, M.J. Burish, J.H. Kaas, Effects of muscimol inactivations of functional domains in motor, premotor, and posterior parietal cortex on complex movements evoked by electrical stimulation, Journal of Neurophysiology. 111 (2014) 1100–1119.

[34] L. Bonazzi, R. Viaro, E. Lodi, R. Canto, C. Bonifazzi, G. Franchi, Complex movement topography and extrinsic space representation in the rat forelimb motor cortex as defined by long-duration intracortical microstimulation, Journal of Neuroscience. 33 (2013) 2097–2107.

[35] A.R. Brown, S. Mitra, G.C. Teskey, J.A. Boychuk, Complex forelimb movements and cortical topography evoked by intracortical microstimulation in male and female mice, Cerebral Cortex. 33 (2023) 1866–1875.

[36] A.R. Brown, G.C. Teskey, Motor cortex is functionally organized as a set of spatially distinct representations for complex movements, J Neurosci. 34 (2014) 13574–13585. 10.1523/JNEUROSCI.2500-14.2014.

[37] A.C. Halley, M.K. Baldwin, D.F. Cooke, M. Englund, L. Krubitzer, Distributed motor control of limb movements in rat motor and somatosensory cortex: the sensorimotor amalgam revisited, Cerebral Cortex. 30 (2020) 6296–6312.

[38] T.C. Harrison, O.G. Ayling, T.H. Murphy, Distinct cortical circuit mechanisms for complex forelimb movement and motor map topography, Neuron. 74 (2012) 397–409.

[39] D. Ramanathan, J.M. Conner, M.H. Tuszynski, A form of motor cortical plasticity that correlates with recovery of function after brain injury, Proceedings of the National Academy of Sciences. 103 (2006) 11370–11375.

[40] M. Budri, E. Lodi, G. Franchi, Sensorimotor restriction affects complex movement topography and reachable space in the rat motor cortex, Frontiers in Systems Neuroscience. 8 (2014) 231.

[41] C.C. Wolsh, R.M. Brown, A.R. Brown, G.A. Pratt III, J.A. Boychuk, Extensive complex neocortical movement topography devolves to simple output following experimental stroke in mice, Frontiers in systems neuroscience. 17 (2023) 1162664.

[42] N.A. Young, J. Vuong, C. Flynn, G.C. Teskey, Optimal parameters for microstimulation derived forelimb movement thresholds and motor maps in rats and mice, Journal of neuroscience methods. 196 (2011) 60–69.

[43] A.R. Brown, M. Martinez, Chronic inactivation of the contralesional hindlimb motor cortex after thoracic spinal cord hemisection impedes locomotor recovery in the rat, Experimental neurology. 343 (2021) 113775.

[44] E. Accili, G. Redaelli, D. DiFrancesco, Differential control of the hyperpolarization-activated current (i (f)) by cAMP gating and phosphatase inhibition in rabbit sino-atrial node myocytes, The Journal of physiology. 500 (1997) 643–651.

[45] M. Castro-Alamancos, J. Borrell, Functional recovery of forelimb response capacity after forelimb primary motor cortex damage in the rat is due to the reorganization of adjacent areas of cortex, Neuroscience. 68 (1995) 793–805.

[46] M.A. Castro-Alamancos, L.M. Garcia-Segura, J. Borrell, Transfer of function to a specific area of the cortex after induced recovery from brain damage, European Journal of Neuroscience. 4 (1992) 853–863.

[47] S.Y. Kim, J.E. Hsu, L.C. Husbands, J.A. Kleim, T.A. Jones, Coordinated plasticity of synapses and astrocytes underlies practice-driven functional vicariation in peri-infarct motor cortex, The Journal of Neuroscience. 38 (2018) 93–107.

[48] R.J. Nudo, B.M. Wise, F. SiFuentes, G.W. Milliken, Neural substrates for the effects of rehabilitative training on motor recovery after ischemic infarct, Science. 272 (1996) 1791–1794.

[49] E. Rouiller, X. Yu, V. Moret, A. Tempini, M. Wiesendanger, F. Liang, Dexterity in adult monkeys following early lesion of the motor cortical hand area: the role of cortex adjacent to the lesion, European Journal of Neuroscience. 10 (1998) 729–740.

[50] M.L. Starkey, C. Bleul, B. Zorner, N.T. Lindau, T. Mueggler, M. Rudin, M.E. Schwab, Back seat driving: hindlimb corticospinal neurons assume forelimb control following ischaemic stroke, Brain. 135 (2012) 3265–3281. 10.1093/brain/aws270.

[51] J.A. Kleim, S. Barbay, N.R. Cooper, T.M. Hogg, C.N. Reidel, M.S. Remple, R.J. Nudo, Motor learning-dependent synaptogenesis is localized to functionally reorganized motor cortex, Neurobiology of learning and memory. 77 (2002) 63–77.

[52] J.A. Kleim, S. Barbay, R.J. Nudo, Functional reorganization of the rat motor cortex following motor skill learning, Journal of Neurophysiology. 80 (1998) 3321–3325.

[53] R.J. Nudo, G. Milliken, W.M. Jenkins, M.M. Merzenich, Use-dependent alterations of movement representations in primary motor cortex of adult squirrel monkeys, Journal of Neuroscience. 16 (1996) 785–807.

[54] R.J. Nudo, G.W. Milliken, Reorganization of movement representations in primary motor cortex following focal ischemic infarcts in adult squirrel monkeys, Journal of neurophysiology. 75 (1996) 2144–2149.

[55] R.M. Glanz, J.C. Dooley, G. Sokoloff, M.S. Blumberg, Sensory coding of limb kinematics in motor cortex across a key developmental transition, The Journal of Neuroscience. 41 (2021) 6905–6918.

[56] A.C. Singleton, A.R. Brown, G.C. Teskey, Development and plasticity of complex movement representations, Journal of neurophysiology. 125 (2021) 628–637.

[57] A.R. Brown, G.M. Coughlin, G.C. Teskey, Seizures alter cortical representations for complex movements, Neuroscience. 449 (2020) 134–146.

[58] A.R. Brown, M. Martinez, Ipsilesional motor cortex plasticity participates in spontaneous hindlimb recovery after lateral hemisection of the thoracic spinal cord in the rat, Journal of Neuroscience. 38 (2018) 9977–9988.

[59] H. Braak, E. Braak, Neuropathological stageing of Alzheimer-related changes, Acta neuropathologica. 82 (1991) 239–259.

[60] M.J. Grothe, H. Barthel, J. Sepulcre, M. Dyrba, O. Sabri, S.J. Teipel, A.s.D.N. Initiative, A.s.D.N. Initiative, In vivo staging of regional amyloid deposition, Neurology. 89 (2017) 2031–2038.

[61] D.A. Cahn-Weiner, J. Grace, B.R. Ott, H.H. Fernandez, J.H. Friedman, Cognitive and behavioral features discriminate between Alzheimer’s and Parkinson’s disease, Cognitive and Behavioral Neurology. 15 (2002) 79–87.

[62] L. Serra, R. Perri, M. Cercignani, B. Spanò, L. Fadda, C. Marra, G.A. Carlesimo, C. Caltagirone, M. Bozzali, Are the behavioral symptoms of Alzheimer’s disease directly associated with neurodegeneration?, Journal of Alzheimer’s Disease. 21 (2010) 627–639.

[63] K. Shinosaki, T. Nishikawa, M. Takeda, Neurobiological basis of behavioral and psychological symptoms in dementia of the Alzheimer type, Psychiatry and Clinical Neurosciences. 54 (2000) 611–620.

[64] B.T. Hyman, G.V. Hoesen, A. Damasio, Memory-related neural systems in Alzheimer’s disease: an anatomic study, Neurology. 40 (1990) 1721–1721.

[65] A. Kargazhanov, R. Aiken, K. Hawkins, R. Lopez, A. Nawaz, G. Srivastava, C. Miller, W. Bogen, C. Long, D. Morgan, Evaluating the peripheral nervous system pathology of Alzheimer’s disease utilizing a functional human NMJ microphysiological system, Alzheimer’s & Dementia. 22 (2026) e71281.

[66] Y.-M. Kuo, T.A. Kokjohn, M.D. Watson, A.S. Woods, R.J. Cotter, L.I. Sue, W.M. Kalback, M.R. Emmerling, T.G. Beach, A.E. Roher, Elevated Aβ42 in skeletal muscle of Alzheimer disease patients suggests peripheral alterations of AβPP metabolism, The American journal of pathology. 156 (2000) 797–805.

[67] Y. Ogawa, Y. Kaneko, T. Sato, S. Shimizu, H. Kanetaka, H. Hanyu, Sarcopenia and muscle functions at various stages of Alzheimer disease, Frontiers in neurology. 9 (2018) 710.

[68] L. Li, Q. Shi, B. Fang, Y. Liu, X. Liu, Y. Shu, Y. Deng, Y. Liu, H. Li, J. Zhou, Functional connectivity changes in primary motor cortex subregions of patients with obstructive sleep Apnea, Brain and Behavior. 15 (2025) e70698.

